# Illuminati, a novel form of gene expression plasticity in *Drosophila* neural stem cells

**DOI:** 10.1101/2021.04.28.441783

**Authors:** Alix Goupil, Jan Peter Heinen, Fabrizio Rossi, Riham Salame, Carole Pennetier, Anthony Simon, Patricia Skorski, Anxela Lauzao, Allison Bardin, Renata Basto, Cayetano Gonzalez

## Abstract

With the aim of developing a genetic instability (GI) sensor in *vivo* we used the well-established Gal80/Gal4-UAS system combined with a visual GFP marker in *Drosophila*. We generated a collection of 25 *Drosophila* lines carrying GAL80 transgenes in different locations in all major chromosomes (X, Y, II, and III). We found low rates of GFP cells in epithelial tissues such as wing discs. In contrast, in larval brains, GFP positive clusters containing neural stem cells- also called neuroblasts (NBs)- and their offspring, were highly frequent. Using genetic and imaging-based approaches, we show that GFP NBs do not result from aneuploidy or mutations in the *GAL80* gene, but rather by stochastic repression of *GAL80* expression. We named this novel type of gene expression instability Illuminati. Importantly, Illuminati frequency is influenced by environmental and stress conditions. Further, we found that once established, Illuminati can be propagated over many cell cycles.

## Introduction

During development, dynamic control between proliferation and differentiation has to be executed in order to reproducibly generate organs and tissues (Rué and Martinez Arias 2015). The establishment of appropriate developmental programs is tightly controlled in space and in time, such as in the optic lobe of the *Drosophila* larval brain. Here the combination of temporal and spatial axes in a set of neural stem cells generates highly complex neuronal diversity (Isshiki et al., 2001; Li et al., 2013; Suzuki et al., 2013; Erclik et al., 2017). Patterns of gene expression can be established by epigenetic modifications in proliferating cells, which can then be inherited by daughter cells and stably maintained over time. Genome plasticity such as the control of gene expression in response to environmental changes also influences tissue and organ behavior with important consequences in organism fitness (Tian and Marsit 2018).

Most cells of a given organism present the same genetic information, which is transmitted throughout generations to maintain genetic stability. Genetic instability (GI), which is used here as a general term to describe chromosome number variations or mutations in DNA leading to chromosome rearrangements defects, cause a variety of diseases including developmental disorders and cancer. Whole genome sequencing (WGS) and single cell analysis have been ground-breaking to identify the frequency and type of GI in health and disease (Knouse et al. 2014; J.-K. Lee et al. 2016). However even if highly informative, WGS does not address GI during a dynamic period of time. Ideally, a GI probe should enable the detection of GI cell birth, while allowing to monitor GI cell behavior or fate in multicellular organisms. Building from previous work using tools to follow chromosome loss or structural chromosome re-arrangements in *Drosophila* embryos, developing somatic cells or adult intestine stem cells (Szabad and Würgler 1987; Carmena et al. 1991; Szabad, Bellen, and Venken 2012; Siudeja et al. 2015), we have conceived a fluorescence-based method to analyze GI in *Drosophila melanogaster*. This system is based on the bipartite Gal4/UAS for detection of green fluorescent protein (GFP) combined with the repressor *GAL80* inserted at specific chromosome sites in all major fly chromosomes (X, Y, II and III). In principle, conditions that remove or inactivate Gal80 in cells that express the bipartite system *GAL4::UAS* combined with a GFP reporter can originate a green fluorescent cell (Figure 1A). The frequency of GFP expressing cells (GFP+) and their lineage have the potential to provide an estimate of the time in development when the initial triggering GI event took place. Analysis of 25 different *GAL80* fly lines corresponding to different chromosome locations revealed only the presence of a low number of GFP+ cells in epithelial tissues such as the wing disc, consistent with low levels of GI in these tissues. Strikingly however, in developing brains a large number of green cells, mostly neural stem cells-also known as neuroblasts (NBs) and their progeny was detected. Using a variety of methods, we show that unexpectedly, these cells do not result from chromosome loss or mutations in *GAL80*. Instead, they seem to result from an instability in gene expression pattern, that we named Illuminati-specific of the *Drosophila* brain, influenced by environmental or stress inducing conditions and maintained over many cell cycle generations.

**Figure 1:**
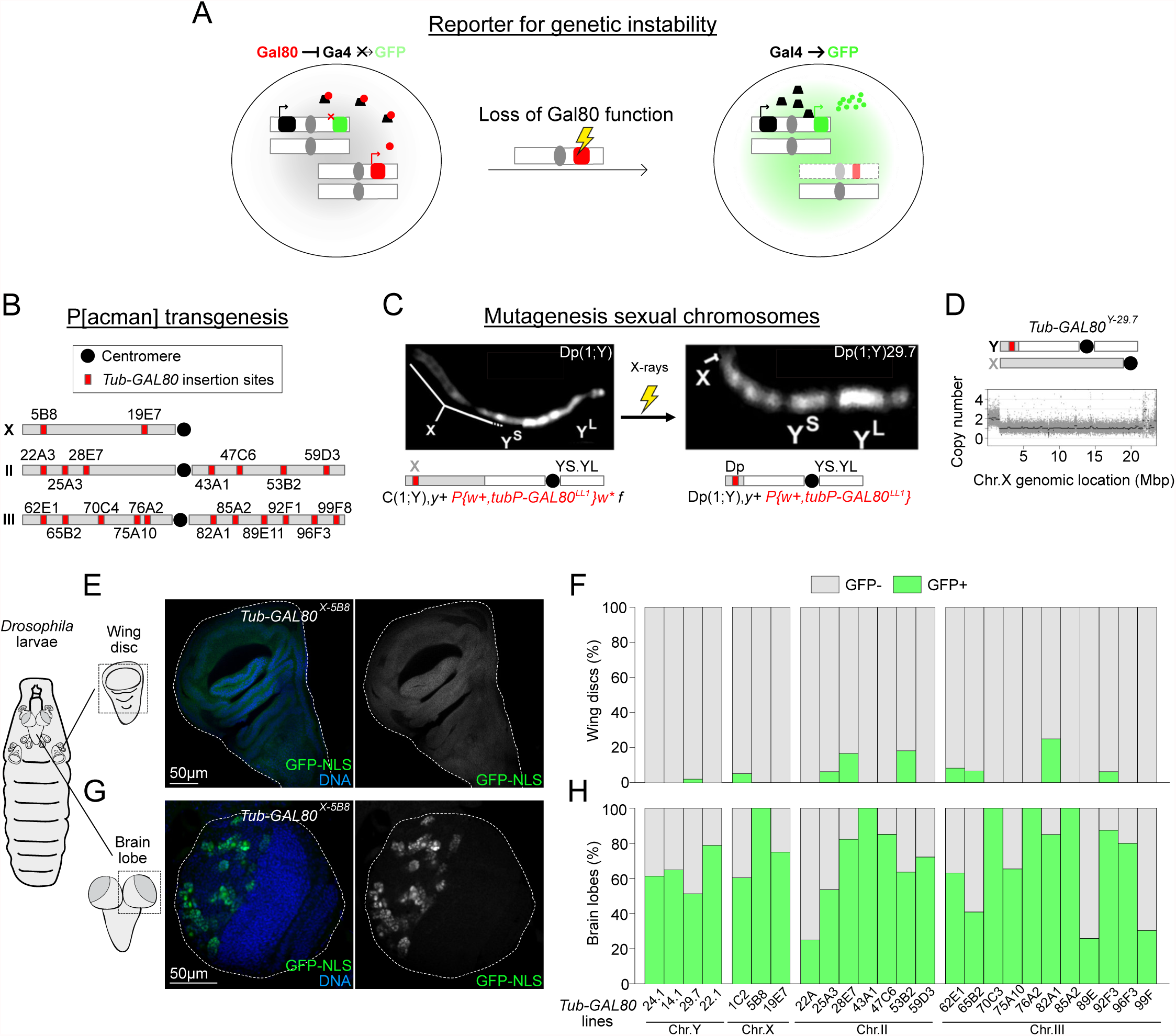
A novel strategy to monitor GI based on the GAL4/GAL80 system. (A) Schematic representation of the genetic system to monitor GI in any *Drosophila* tissue. On the left-the presence of Gal80 inhibits Gal4 and so represses GFP expression, cells are black. On the right-upon loss of Gal80 functions due to GI (aneuploidy or mutations that impair *GAL80* expression) Gal4 is released from Gal80 repression and promotes GFP expression, and so cells are green. (B) Representative map of the *Tub-GAL80* insertion sites on *Drosophila* chromosomes X, II and III obtained by P[acman] transgenesis. (C) Strategy to generate a duplication of the distal part of the X to the Y chromosome carrying a *Tub-GAL80* insertion using recombination followed by X-rays mutagenesis. Chromosomes are represented with DAPI labelling and schematized below images. (D) Graph showing the X chromosome copy number from sequencing of *Tub-GAL80^Y-29.7^* males. Sexual chromosomes are schematized above the graph: the*Tub-GAL80* cassette (red square) is located on a small part of the X chromosome (grey) duplicated to the Y chromosome (white). (E and G) Schematic representation of the *Drosophila* L3 larvae and images of wing disc (E) and brain lobe (G) for *Tub-GAL80^X-5B8^* condition. Whole mount tissues were labeled for GFP (green and grey) and with DAPI for DNA (blue). White dotted lines delimitate tissues. (F and H) Graph bars summarizing the percentage of wing discs (F) and brain lobes (H) without (grey) and with GFP signal (green) from the screen of all *Drosophila* lines carrying one copy of the *Tub-GAL80* cassette on chromosomes X, Y, II and III (n=7 to 20 wing discs/condition and n=14 to 88 BLs/condition).

## Results

### A novel strategy to monitor genetic instability based on the Gal4/Gal80 system

We sought to generate tools to determine the background level of genetic instability (GI) in different cell lineages during development in *Drosophila*. To this end we made use of the repressive effect of Gal80 on the Gal4::UAS transcription activation system. Flies that carry a *UAS-GFP* transgene and express both *GAL4* and *GAL80* from constitutive promoters cannot express GFP because Gal80 prevents Gal4 from binding to UAS (T. Lee and Luo 1999). GI affecting Gal80 function will therefore switch on GFP expression resulting in green cells that will include the original cell and its offspring. Such a reporter could in principle be used to analyze the frequency and to time the onset of GI in any tissue (Figure 1A).

To this end we generated a collection of *Drosophila* strains carrying *GAL80* insertions in all four major chromosomes (X, Y, II and III). To generate the lines carrying *GAL80* on the X, II and III chromosomes we designed a new vector carrying a *GAL80* version optimized for *Drosophila* codon usage (Pfeiffer et al. 2010). This sequence is under the control of the ubiquitous Tubulin 1α promotor – *Tub-GAL80* (O’Donnell, Chen, and Wensink 1994; T. Lee and Luo 1999). To minimize the risk of undesired positional effects that could affect *GAL80* expression, we generated these lines using the targeted insertion ΦC31-recombination attB P[acman] system, which has been previously established and validated (Venken et al. 2006). A total of 20 *Drosophila* transgenic lines, each carrying one copy of the *Tub-GAL80* transgene inserted at different genomic regions was obtained (Figure 1B). Each of these lines is referred to as *Tub-GAL80* followed by the designation of the chromosome and insertion site in superscript (e.g *Tub-GAL80^X-5B8^*-*GAL80* inserted on the X chromosome at location 5B8).

To generate *GAL80* insertions on the Y chromosome we first recombined the *P{w+, tubP-GAL80LL1}* transgene located distally on the X chromosome (T. Lee and Luo 1999) into a *C(1;Y)* chromosome that carries a fully functional fusion between the X and Y chromosomes sharing a single centromere. The resulting *C(1;Y) P{w+, tubP-GAL80LL1}* was then subjected to x-ray mutagenesis to generate large deletions that remove most of the X chromosome leaving only the distal most part of the X chromosome of *C(1;Y)* attached to a fully functional Y chromosome; i.e. transforming the original *C(1;Y) P{w+, tubP-Gal80LL1}* in a Dp(1;Y) *P{w+, tubP-GAL80^LL1^}* (Cook et al. 2010). From a total of 38.206 *C(1;Y) P{w+, tubP-GAL80^LL1^}* chromosomes we obtained four different lines that will be referred to as *Tub-GAL80^Y^* followed by a number to identify each line. Banding of mitotic chromosomes labeled with DAPI revealed the corresponding *GAL80-Y* as a short euchromatic region attached distally to the Y heterochromatin (Figure 1C). Genome wide sequencing confirmed these as X chromosome distal duplications and identified the exact break point. The size of these duplications ranges from 1.08Mbp to 1.66Mbp (Figure 1D).

To determine the suitability of our collection of GI reporters we quantified the basal rate of GFP expression in wing imaginal discs and larval brains. As expected, we found that regardless of the chromosome where the *GAL80* transgene is inserted (X, Y, II or III), the vast majority of wing discs did not contain GFP expressing cells (GFP-) (416/429 in total) (Figure 1E-F and compare with Supplementary Figure 1A-F for controls). In 10 out of the 13 wing discs that contained GFP positive cells (GFP+), these were restricted to only a few cells within the whole GFP-tissue (10 wing discs from 7 different *Tub-GAL80* constructs). Interestingly, only one single *Tub-GAL80* line-*TubGAL80^III-82A1^*-had a high number of GFP+ cells, and this only occurred in 3 out of 12 wing discs analyzed (Supplementary Figure 1G-H).

In stark contrast to discs, most larval brains examined presented clusters of GFP+ cells at rates that varied substantially, but were always at least one order of magnitude greater than that observed in wing discs (Figure 1G-H and compare with Supplementary Figure 1A-F for controls).

We considered the possibility that differences between the number of GFP clones between wing discs and brains might be explained by differences in cell number and mitotic activity between the two tissues. However, using the MARCM system - heat-inducible recombination of FRT sites by FLP - to induce wild type clones during mitosis (Supplementary Figure 2A), we only observed very minor differences in the frequency of GFP+ cell clusters in wing discs (mean= 22.4, n= 7) and brain lobes (mean= 16.6, n=10) (Supplementary Figure 2B-D). Thus, differences in cell number and mitotic activity cannot account for the high frequency of GFP+ cells in the brain, when compared to wing discs.

Our results show that, as expected, the large majority of the *Tub-GAL80* insertions generated in this study efficiently repress Gal4*::UAS-GFP* expression in the larval wing disc epithelium. They also show that GI is a very rare event in this tissue. Moreover, our results also reveal an unexpected number of GFP+ cells in the brain that is not dependent on the site of *GAL80* chromosome insertion or on the method used to generate these lines (P-element based or site directed recombination). We thus decided to analyze these cells in more detail.

### The majority of GFP+ clusters comprise central brain neuroblasts and their lineage

The larval brain lobe can be divided in two main regions, the central brain and the optic lobe (Figure 2A). The central brain is composed of neural stem cells, also called neuroblasts (NBs) that divide asymmetrically to self-renew and give rise to smaller and more committed progenitors, the ganglion mother cells (GMCs) (Bello et al. 2008; Boone and Doe 2008; Homem and Knoblich 2012). The optic lobe comprises two proliferative centers, the inner and the outer, which correspond to a pseudo-stratified epithelium called the neuroepithelium (NE). NE cells give rise to neurons necessary for the development of the visual system of the fly. Further, perineural and sub-perineural glial cells with large nuclei are found at the superficial layer of the brain (Pereanu, Shy, and Hartenstein 2005). All these cell types are easily distinguishable by their morphology, position in the brain and expression markers. For instance, the signals Dpn^+^, Dpn^-^/Propero^weak^, Elav^+^, and Repo^+^ label NBs, GMCs, neurons and glial cells respectively. The NE is distinguished by the morphology of its composing cells revealed with actin (Rujano et al. 2013).

**Figure 2:**
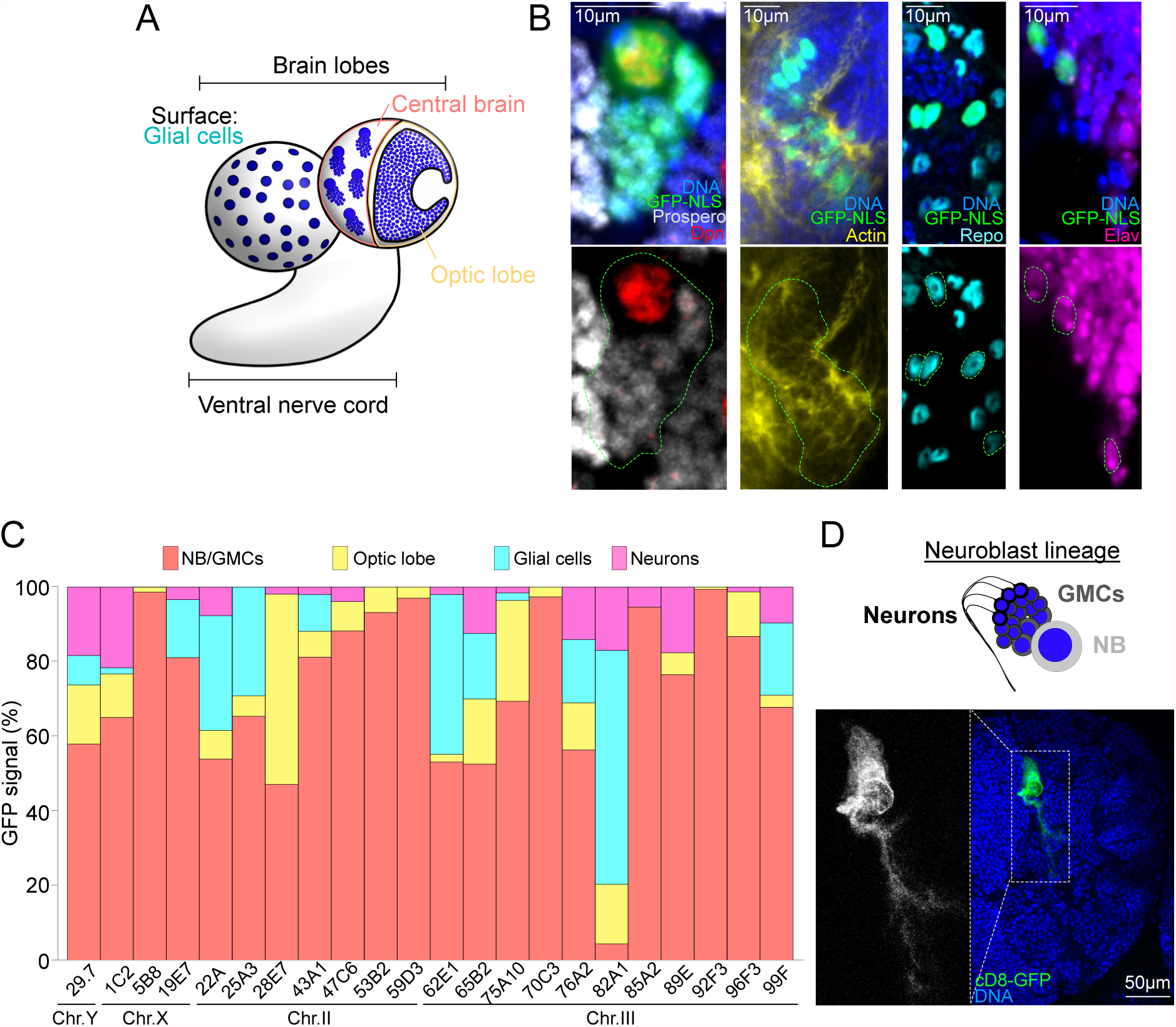
The majority of GFP positive clusters correspond to NB lineages in larval brains. (A) Schematic representation of the *Drosophila* larval brain and its different cell types. Glial cells with large nuclei are present at the surface of the brain lobes. The core of the brain lobe is divided in two parts, the central brain and the optic lobe. The central brain is composed of NBs that divide asymmetrically to generate GMCs that will then give rise to neurons. The optic lobe is composed of neuroepithelial (NE) cells that divide symmetrically. (B) Zoom inset images in whole mount brain lobes labeled for GFP (green) and for specific markers of the different cell types: Dpn^+^ NBs (red) with Prospero^weak^ GMCs (grey), cells of the NE from the optic lobes distinguishable by the specific F-actin organization (yellow), Repo^+^ glial cells (cyan blue) and individual Elav^+^ neurons (pink). DNA is shown in blue. Green dotted lines surround GFP+ clusters or cells. (C) Graph bar showing the percentage of cells showing GFP+ signals for each cell type of the brain lobe: NB/GMCs clusters (red), optic lobe cells (yellow), individual glial cells (cyan blue) and individual neurons (pink) (n=7 to 43 BLs/condition). (D) Image of whole mount brain and zoom inset of GFP+ cluster containing a NB and a full lineage including neurons in larvae expressing the membrane marker *UAS-cD8-GFP* in addition to *GAL4* and *Tub-GAL80^Y-29.7^*. Neuroblast lineage is schematized above the image.

Taking advantage of this wealth of markers we determined the identity of the GFP+ cells that appeared in *GAL80,GAL4::UAS-GFP* brains. Analysis of the 22 fly lines comprising 590 brain lobes from all the X, Y, II and III chromosome *Tub-GAL80* lines (minimum of 14 brain lobes per *Tub-GAL80* insertion line) revealed that the number and identity of GFP+ cells varied between different *Tub-GAL80* lines (Figure 2B-C). Three extreme examples are *Tub-GAL80^X-5B8^* that presented a high number of GFP+ cells most of which were NBs and associated GMCs, very few cells in the optic lobe, and neither neurons or glial cells; *Tub-GAL80^III-82A1^* that also showed a high number of GFP+ cells, mostly neurons and glia and few NBs/GMCs clusters; and *Tub-GAL80^II-22A^* where green cells in each category were rare. However, notwithstanding inter-line variability, plotting the mean frequency for each cell type for all the *Tub-GAL80* lines showed that most GFP+ cells were located in the central brain and most of them presented NBs and associated GMCs (Figure 2C). In addition, in larvae expressing a membrane UAS-GFP (*UAS-cD8::GFP*), which allows to visualize fluorescence in cell processes, we frequently found GFP+ clusters that contained one NB and its lineage including neurons (Figure 2D). Importantly, in the MARCM experiments described above (Supplementary Figure 2A), clones were more frequently in the optic lobe (58%) than in NBs and associated GMCs from the central brain (13%) (Supplementary Figure 2E). These results illustrate once more that differences in mitotic activity cannot account for the predominance of GFP+ in central brain NBs. It is important to mention that we did not find a trend in the position or spatial arrangement of GFP+ NBs or even other cell types within the different brain lobes analyzed for each *Tub-GAL80* insertion. These observations suggest that cells expressing GFP are randomly positioned in the central brain.

Altogether these results demonstrate that although the frequency and cell types do vary among different *GAL80* lines, most GFP+ cells in the brain represent NBs and their lineage localized in the central brain region.

### Analysis of *Tub-GAL80* lines alerts its use for MARCM analysis

The Gal4/Gal80 system is widely and routinely used by the *Drosophila* community to control gene expression. Importantly, this system has been used in MARCM experiments for neuronal lineage tracing in the developing *Drosophila* brain (T. Lee and Luo 1999; Ren et al. 2016). This is based on the loss of heterozygosity (LOH) after mitotic recombination by the heat-induced FLP recombinase at specific FRT sites. LOH generates labeled mutant clones that lack the *Tub-GAL80* sequence and unlabeled wild type cells (WT) homozygous for *Tub-GAL80* (Supplementary Figure 3A). Our results predict that at a fraction of GFP+ cells might not result from LOH. To test this possibility, we induced MARCM *sas4* mutant (*sas4^mut^*) clones by crossing heat-shock *FLP; FRT82B sas4^mut^* flies with *UAS-GFP-NLS; Tub-GAL4, FRT82B, Tub-GAL80* from the Bloomington stock center BLSC (#5132) (T. Lee and Luo 1999). The *sas4* gene encodes for a protein essential for centriole duplication and *sas4^mut^* cells lack centrosomes (Basto et al. 2006). Larvae were heat-shocked at 37°C for 1H to induce FLP mediated recombination. We analyzed GFP+ clones in L3 brains and wing discs. As expected, we observed GFP+ *sas4^mut^* clones without centrosomes (absence of Sas4 and Cnn-a centrosome component) (Supplementary Figure 3B). However, as we expected, we also observed GFP+ clones that contained centrosomes (Supplementary Figure 3C). These results indicate that the WT *sas4* gene was present, most likely because these latter clones were not generated through *Tub-GAL80* sequence loss upon FLP/FRT-mediated LOH. Also, as expected GFP+ clones were also observed in control brains that were not heat-shocked (Supplementary Figure 3E-F). Importantly, in contrast to brains, GFP+ clones in wing discs were only observed after heat-shock induction and these clones did not express *sas4* (Supplementary Figure 3D and 3G).

**Figure 3:**
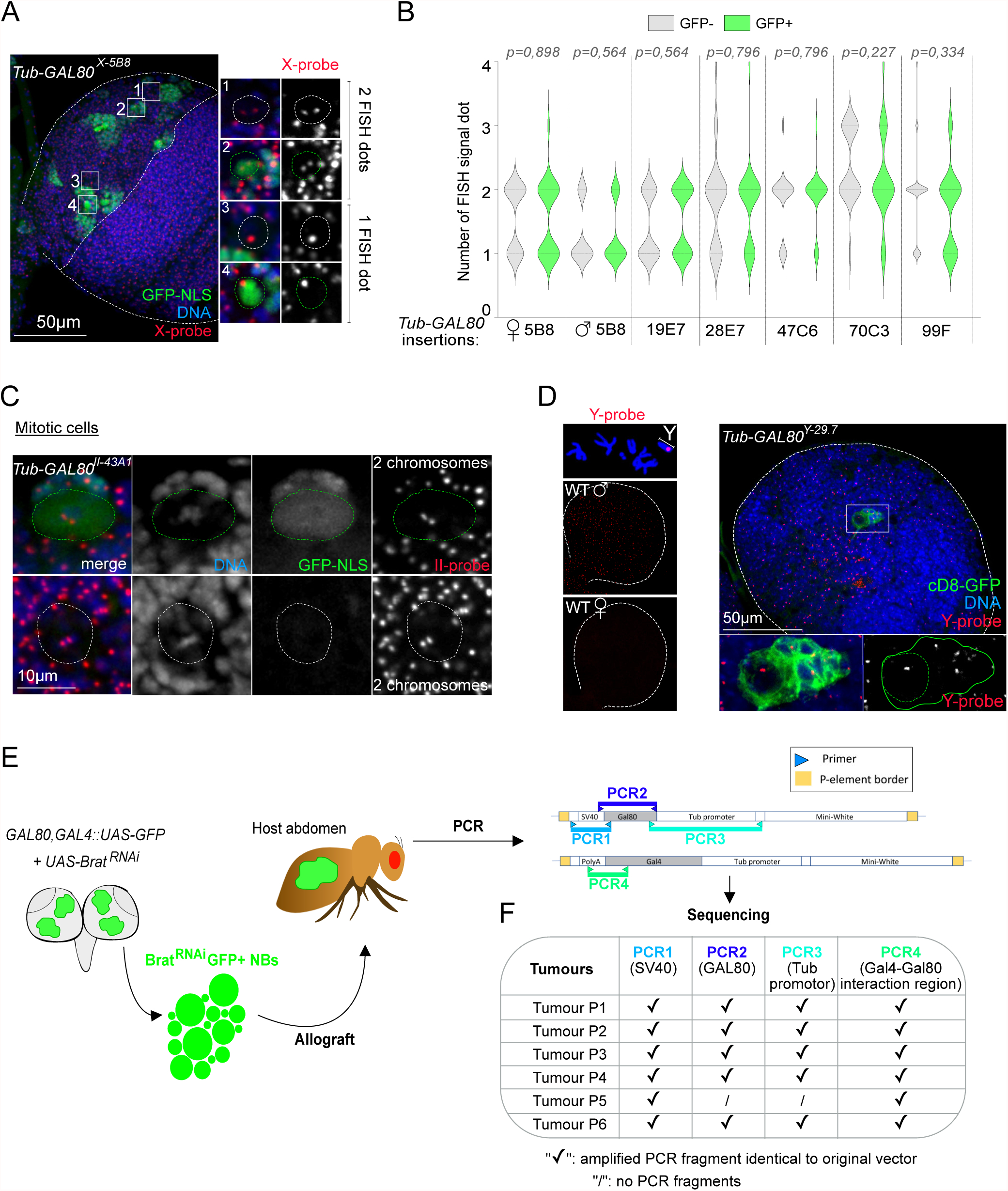
GFP positive cells are not a bi-product of aneuploidy or GI in the larval brain. (A and C-D) Fluorescent *in situ* hybridization of whole mount brains using probes for chromosomes X (A), II (C) and Y (D) (red and grey) combined with GFP labeling (grey and/or green) and DAPI for DNA (grey and/or blue). White dotted lines surround brain lobes and/or GFP^_^ NBs. Green continuous and dotted lines surround GFP+ clusters and cells, respectively. (B) Violin plot representing the number of FISH signal dots between GFP+ and GFP^_^ cells. FISH signals correspond to the chromosomes X, II or III for conditions where *Tub-GAL80* was inserted at positions 5B8 (n=43 cells for females and n=50 cells for males) and 19E7 (n=47 cells), 28E7 (n=50 cells) and 47C6 (n=50 cells) or 70C3 (n=47 cells) and 99F (n=60 cells), respectively. FISH signals are variable between conditions but similar between GFP+ (green) and GFP^_^ (grey) cells from the same condition. Statistical non-significance was determined by a Mann-Whitney test and p corresponds to the p-value. (C) GFP+ and GFP^_^ NBs are diploid as they present two dots for the two chromosomes II. (D) Confirmation of Y-probes efficiency for Y chromosome detection on metaphase spread chromosomes and whole mount brains in WT males (XY). As expected, no signal was detectable in WT females. In males containing the Gal4/Gal80/GFP system, GFP+ clusters are not aneuploid as shown by the detection of the *GAL80*-containing Y chromosome by Y-probes (n=33 NBs). (E) Schematic representation of the protocol used to sequence the *GAL80* and *GAL4* genes in GFP+ brain cells. *UAS-Brat^RNAi^* was used to induce tumors and thus, drastic increase of the GFP+ population to obtain sufficient DNA. *Tub-GAL80* coding and regulatory sequences were amplified by PCR and subsequently sequenced. (F) Table summarizing the results obtained for each tumor line (n= 6 tumor samples).

Our results show the presence of false positive cells using the MARCM system in the larval brain, but not in the wing disc.

### GI does not account for the unexpected GFP+ cells in the brain

Two mechanistically different processes can in principle account for the appearance of GFP+ cells in the brain: aneuploidy i.e loss of the chromosome that carries the *GAL80* transgene, or DNA damage leading to *Gal80* mutations leading to lack of expression or to expression of inactive mutant forms of Gal80.

To investigate if GFP+ cells were aneuploid, we performed fluorescent in situ hybridization (FISH) using probes generated against the X, II and III chromosomes in brains containing GFP+ cells. Precise ploidy quantification is not simple because the number of FISH positive dots in diploid cells can range from 1 to 4 depending on both cell-cycle stage and extent of chromosome pairing (Joyce et al. 2012). However, after the analysis of the distribution of FISH signals, we conclude that these were similar between GFP+ and GFP-cells (n=347 cells), (Figure 3A-B). Moreover, FISH in mitotic NBs, where FISH signals can be assigned to individual chromosomes (Carmena et al. 1993; Gatti, Bonaccorsi, and Pimpinelli 1994) confirmed that GFP+ cells were not aneuploid (Figure 3C). To unequivocally test whether GFP+ cells result from chromosome loss, we used a probe against the Y-specific satellite AATAC (Figure 3D) (Bonaccorsi and Lohe 1991) in *Tub-GAL80^Y-29.1^* brains. We found that all GFP+ clones (n=33) presented a Y-specific FISH signal that could not be told apart from that of the neighboring cells that did not express GFP. These results show that the high frequency of green cells observed across different *Tub-GAL80* lines cannot be explained by the loss of the *GAL80* bearing chromosome.

We then investigated the possible contribution of GI to the presence of green cells in the brain. We decided to sequence the *Tub-GAL80* transgene to ascertain if mutations could account for loss of Gal80 function. This is technically challenging because GFP+ cells are orders of magnitude less abundant than the surrounding cells that do not express GFP. To circumvent this problem we generated flies that in addition to the usual combination of *GAL80, GAL4::UAS-GFP* transgenes also carried a fourth transgene encoding *UAS-Brat-RNA interference* (*Brat^RNA^*^i^). We reasoned that if the loss of Gal80 function that leads to transcription of the *UAS-GFP* transgene would lead to the concomitant transcription of *UAS-Brat^RNAi^*, certain GFP+ cells could develop as tumors that could be cultured in allografts. This in principle would make it possible to isolate large quantities of DNA from GFP+ cells (Figure 3E). Using this approach, we obtained 6 tumors from which we sequenced the corresponding *Tub-GAL80* regulatory and coding sequences. We also amplified and sequenced the carboxy-terminal *GAL80* binding site of *GAL4* because mutations or deletions in this region result in constitutively active Gal4 in the presence of Gal80 (Ma and Ptashne 1987). We found that in 5 out of 6 tumor samples, the *Tub-promoter, GAL80* CDS and *SV40 terminator* sequences (i.e PCR3, 2, and 1, respectively) were identical to those contained in the original transgene *P{w^+mC^, tubP-GAL80}LL1*. In the remaining sample, PCR failed to amplify fragments PCR2 and PCR3. In all six tumor samples the sequence corresponding to the Gal80 binding site of Gal4 was found to be wild type (Figure 3F).

These results strongly suggest that the vast majority of GFP+ cells observed in *GAL80,GAL4::UAS* larval brains represent a type of functional instability that is not caused by chromosome loss or GI. We will henceforth refer to this unknown phenomenon of “illumination” of NBs in the brain as Illuminati.

### Stoichiometry imbalance contributes to illuminati

To get further insight into the molecular mechanism of Illuminati we studied the consequences of changing the 1:1:1 ratio of Gal80/Gal4/UAS. We first determined the effect of increasing the number of *Tub-GAL80* transgenes. To this end we generated 5 different *Drosophila* recombinant lines harboring two copies of *Tub-GAL80* inserted at distant loci of the same chromosome (1 line for the X chromosome and 2 lines for the II and III chromosomes-referred to as *2xTub-GAL80*). We found that Illuminati was fully suppressed in all brain lobes from two lines, while in the remaining 3 lines, illuminati cells were still detected (Figure 4A and 4C). Interestingly, comparison of females with either *2xTub-GAL80* or *4xTub-GAL80* copies on the X chromosome showed a marked reduction in Illuminati frequency.

**Figure 4:**
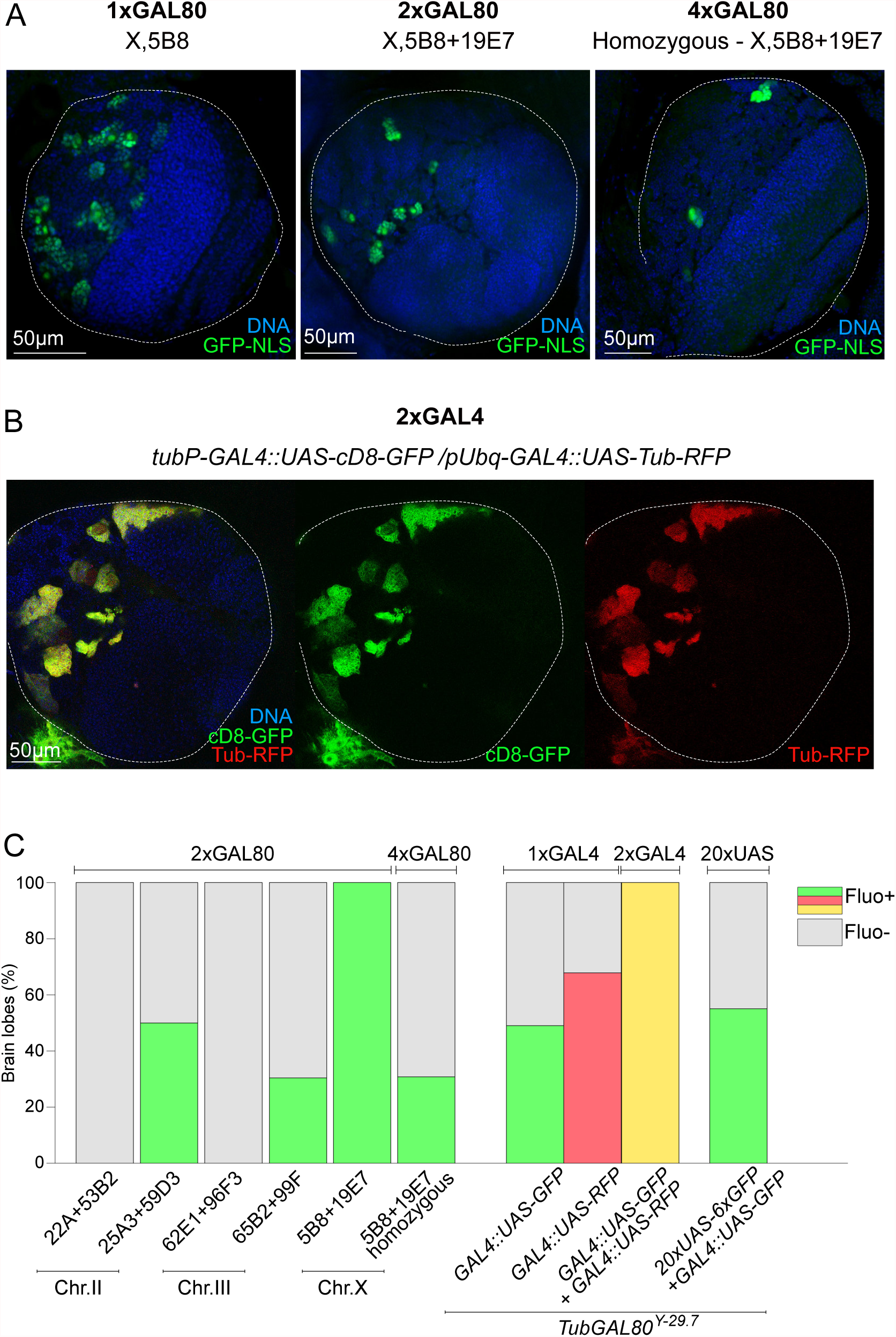
Gal4/Gal80 stoichiometry contributes to the presence of illuminati cells in the larval brain. (A-B) Images of whole mount brain lobes showing DNA (blue), (A) GFP labelling (green) and (B) RFP labelling (red). White dotted lines delimitate brain lobes. (C) Graph bar showing the percentage of brain lobes without (grey) and with fluorescent signal (green, red or yellow) (n=8 to 51 BLs/condition).

Consistent with the inhibitory effect of several *GAL80* transgenes, Illuminati was notably enhanced in flies that carried two *GAL4* transgenes (Figure 4B-C). This increase in Illuminati frequency was largely accounted for by clones in the central brain (clone frequency was not increased in optic lobe) containing NBs and associated GMCs. Interestingly, Illuminati clones in individuals carrying two *GAL4* transgenes presented a wide range of fluorescence intensity, i.e fluorescence signal within a given clone remains remarkably stable but it is rather variable from clone to clone. Illuminati clones in individuals carrying these two *GAL4* together with 2 UAS sequences, each driving a different fluorescent protein (*UAS-GFP* and *UAS-RFP*) co-expressed both colors at roughly similar fluorescence intensities (i.e. clones that present weak and strong GFP signal also present weak and strong RFP signal, respectively) (Figure 4B). The presence of one additional UAS transgene that carried a tandem repeat of 20X UAS had no effect on Illuminati frequency (Figure 4C).

Altogether, these data are consistent with a model in which Illuminati cells arise because of Gal80 levels stochastically falling below the critical concentration threshold that is required to efficiently suppress Gal4::UAS driven transcription. Different fluorescent intensities may reflect a dynamic range of Gal80 function levels, all below the threshold, but still able to partially inhibit the Gal4::UAS system to a greater or lesser extent. Adding an extra *GAL4* transgene raises the threshold, hence increasing the number of cells in which Gal80 function falls below it, which indeed is reduced by one additional *GAL80* transgene.

### Illuminati NBs maintain a pattern of GFP expression that correlates with lack of *GAL80* expression

To obtain a dynamic view of illuminati we performed long term time-lapse microscopy covering nearly two thirds of the total proliferative window of the central nervous system during third instar larvae. We analyzed 17 *2xTub-GAL80^X-5B8,19E7^* brain lobes with a total number of 167 GFP+ events, all of which restricted to the central brain NB lineages. Consistent with immunofluorescence data from fixed samples, GFP+ clusters corresponded to NBs and their lineage. Regarding GFP+ NBs, we found that the vast majority (94.6%, n=158 out of 167) produced GFP+ GMC cells and maintained GFP+ signals through successive rounds of cell division (Figure 5A-B). Interestingly, minor behaviors were also identified. In 7 NBs (4.2%), the initial GFP+ signal disappeared and the NB became GFP^−^. Further, *de novo* GFP appearance could only be identified in 2 NBs (1.2%), (Figure 5B and Supplementary Figure 4A-B).

**Figure 5:**
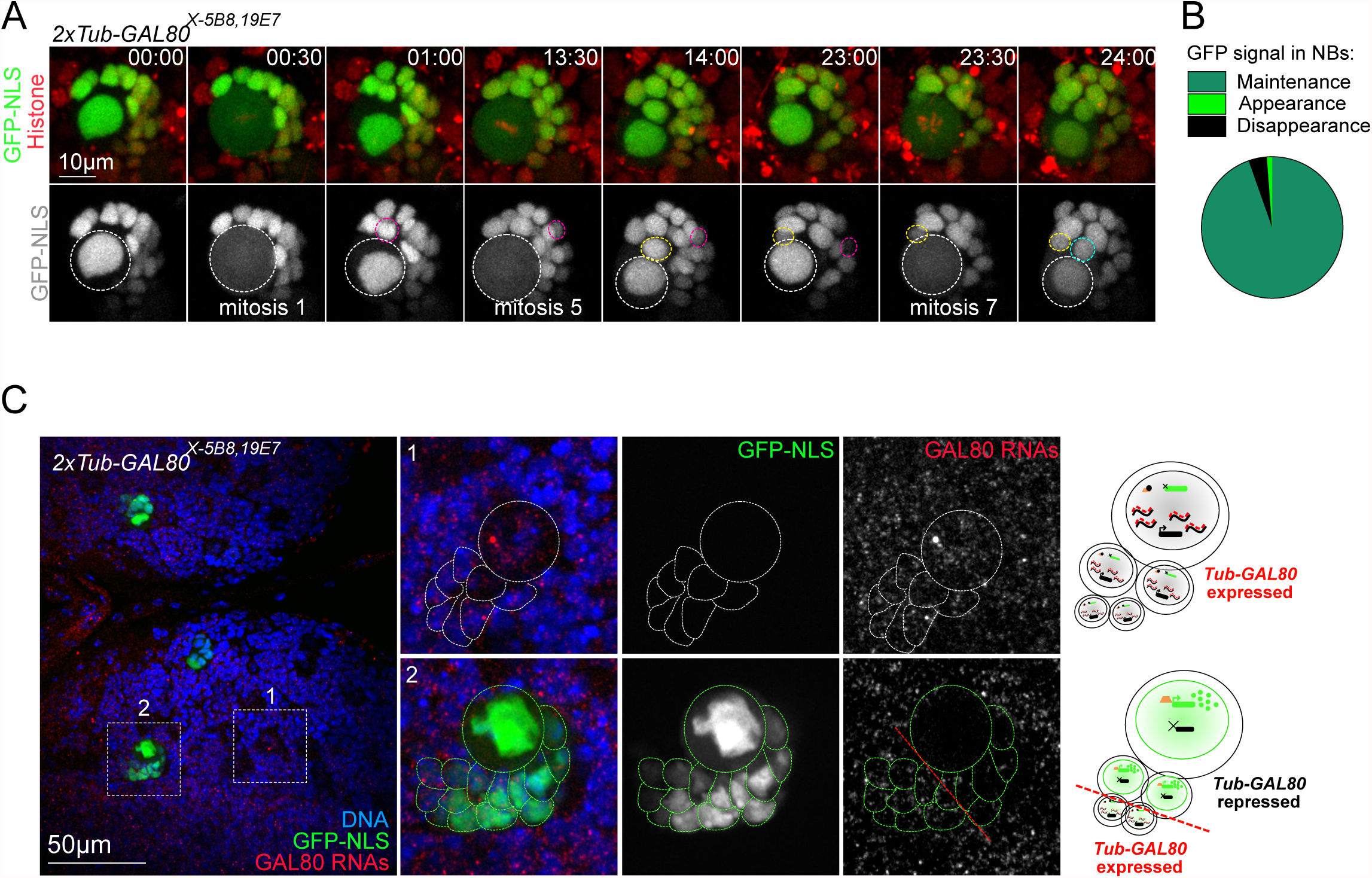
GFP signal in illuminati NBs is maintained throughout several consecutive divisions and correlates with lack of *GAL80* expression. (A) Stills of time-lapse movies of mitotic NBs expressing *2xTub-GAL80^X-5B8,19E7^*, *GAL4::GFP-NLS* (green) and histone-RFP (red) to monitor GFP and chromosome dynamics. White and colored dotted circles surround NBs and daughter GMCs, respectively. (B) Pie chart of the different GFP dynamics in illuminati NBs (n=167 NBs from 17 brain lobes): maintenance and clonal expansion (dark green), appearance (light green) or disappearance (black) of the GFP signal. (C) Images of whole mount brain lobes from RNA FISH experiments with probes against the GAL80 RNAs (red and grey in zoom insets) and labeled with GFP booster (green and grey in zoom insets) and DAPI for DNA (blue). Schematic representation of cells is shown next to the images. White and green dotted lines surround GFP^_^ and GFP+ NB/GMCs clusters, respectively.

Regarding the major category, GFP+ NBs and their progeny, the intensity of the green fluorescence was decreased in GMCs positioned furthest away from the NB. This observation suggests that *GAL80* expression might have been re-established in these cells. To test this possibility, we designed FISH probes that recognize GAL80 mRNAs. We found that GFP+ NBs lack GAL80 RNAs FISH signals that can be easily recognized in GFP^_^ cells. Notably, GAL80 RNAs FISH signal was also detectable in the cells within the GFP+ NB-progeny clusters that present a low fluorescence intensity (Figure 5C).

The results above show that although the probability of illuminati taking place in a NB at any given cell cycle is relatively low, once Illuminati occurs, the large majority of NBs retain the Illuminati status passing out the condition to their offspring through successive cell cycles.

### Illuminati does not result from Position effect variegation or G4::UAS variegation

Illuminati clones reveal an unexpected level of GFP expression variegation. There are two well characterized types of gene expression variegation in *Drosophila*: position effect variegation (PEV) and Gal4::UAS variegation. PEV is caused by the silencing of a gene in certain cells through its proximity to heterochromatin (Elgin and Reuter 2013). Gal4::UAS variegation, on the other hand, refers to the variegated expression of Gal4-driven UAS-genes that can be observed in *GAL4::UAS-EGFP* expressing follicle epithelia. In this tissue, patches of cells containing different fluorescence intensity can be distinguished (M.-C. Lee, Skora, and Spradling 2017; Skora and Spradling 2010).

To assess the possible contribution of PEV or Gal4::UAS variegation to Illuminati, we quantified the frequency of illuminati clones in flies that were heterozygous for each of the following modifiers of variegation: Poly-(ADP-ribose) polymerase (*PARP*), Suppressor of variegation 3-3 (Su(var)3-3, a.k.a Lsd1), *CoRest*, *Six4* and *mutagen-sensitive* 312 (*mus*312) (Tulin, Stewart, and Spradling 2002; Skora and Spradling 2010; M.-C. Lee and Spradling 2014). *PARP* is a strong enhancer of Gal4::UAS variegation, while *Su(var)3-3*, *CoRest*, *Six4*, and *mus312* are suppressors of Gal4::UAS variegation. In addition, Su(var)3-3 is also a strong suppressor of PEV. We found that the frequency of illuminati clones was only slightly, but significantly increased in *PARP*/+ and decreased in *Su(var)3-3*/+ and in *mus312*/+ larvae. In contrast, heterozygous condition for either *Six4* or *mus312* had no significant effect in illuminati frequency (Supplementary Figure 5). Notably, the increase of Illuminati frequency observed in PARP/+ individuals was largely accounted for by NB-containing clones in the central brain.

These results show that like PEV and Gal4::UAS variegation, illuminati is dependent upon the chromatin state. However, the lack of effect of six4 and mus312, together with the quantitatively minor effect of PARP, Su(var)3-3, and CoRest suggests significant mechanistic differences may apply.

### Investigating the sensitivity of Illuminati to stress and environmental conditions

Our results strongly suggest that Illuminati reflect a yet unidentified mechanism by which Gal80 function falls below the minimum level required for efficient inhibition of Gal4::UAS-driven transcription. If that was the case, one could expect Illuminati frequency to be sensitive to experimental conditions that impose a certain level of stress and unset gene expression. To test this hypothesis, we assessed the effect of food composition and temperature.

Using the *2xTub-GAL80^X-5B8,19E7^* line which presents a relatively high frequency of illuminati under normal culturing conditions, we found that a protein-poor medium made of cornmeal and low yeast content, significantly reduces the number of GFP+ clusters when compared to that of flies cultured in standard rich medium (Figure 6A). We next tested the effect of temperature on Illuminati frequency in the same *2xTub-GAL80^X-5B8,19E7^* line. We found that illuminati frequency steadily decreased as culture temperature raises from 18°C to 29°C (Figure 6B). We then tested the effect of temperature on the *Tub-GAL80^Y-29.7^* line that presents a much lower frequency of Illuminati than *2xTub-GAL80^X-5B8,19E7^* line at 18°C. Remarkably, we found that in this line, the Illuminati frequency was strongly enhanced at higher temperatures (Figure 6C). These results put in evidence that illuminati is sensitive to environmental stimuli and reveal that different *GAL80* insertions can respond differently to such stimuli.

**Figure 6:**
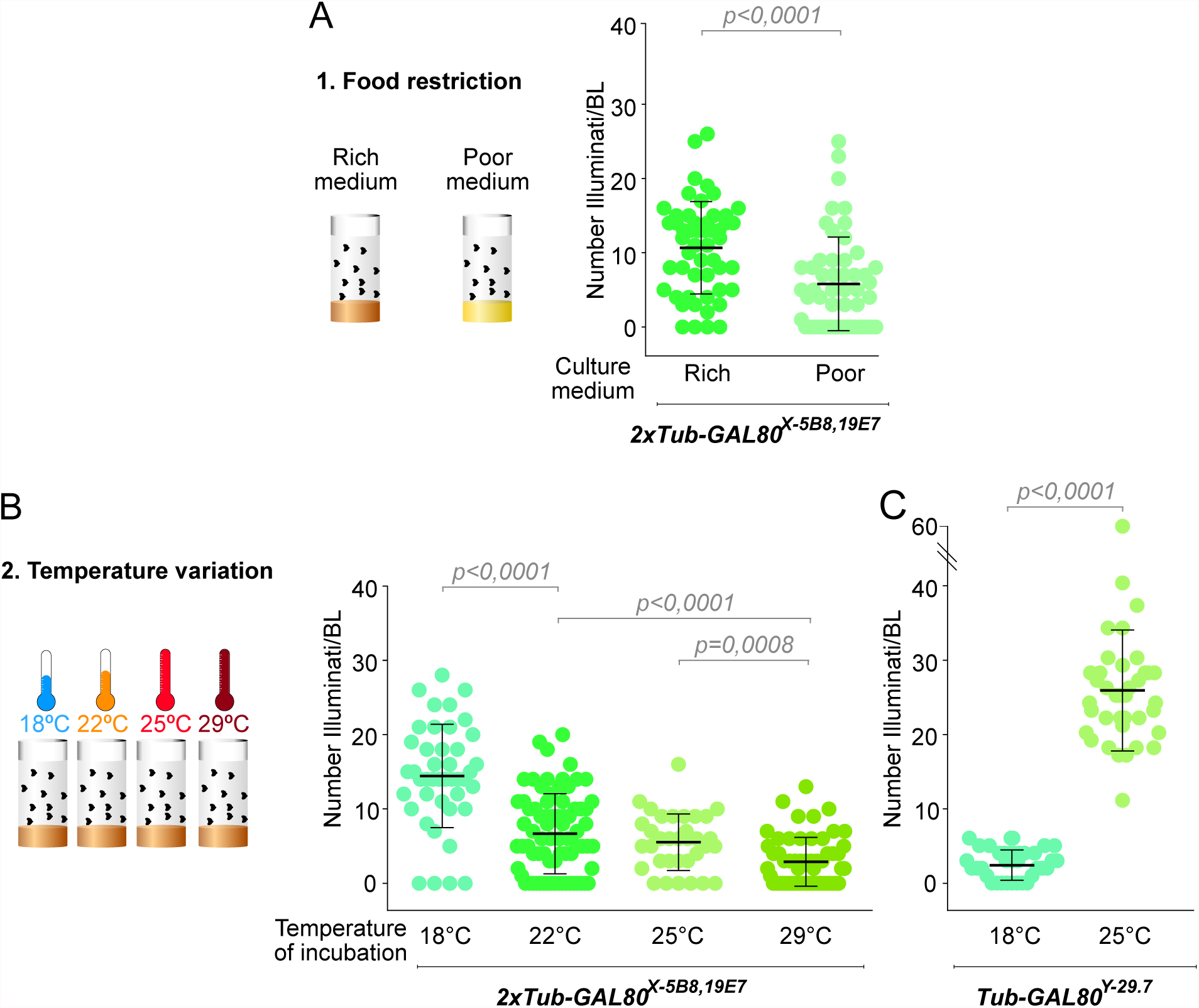
Illuminati is influenced by stress conditions. (A-C) Dot plot showing the number of Illuminati/brain lobe in larvae (A) raised on different culture media and (B-C) at different temperatures. (A) *2xTub-GAL80^X-5B8,19E7^* flies were raised on protein-rich (n=70 BLs from 35 brains) or -poor medium (n=62 BLs from 31 brains). (B) *2xTub-GAL80^X-5B8,19E7^* flies were raised at 18°C (n=43 BLs from 22 brains), 22°C (n=76 BLs from 38 brains), 25°C (n=34 BLs from 17 brains) or 29°C (n=58 BLs from 29 brains). (C) *Tub-GAL80^Y-29.7^* flies were raised at 18°C (n=42 BLs from 21 brains) or 25°C (n=39 BLs from 20 brains). Statistical significance was determined by a Mann-Whitney test and p corresponds to the p-value. Error bars correspond to the means ± SD.

### Illuminati can contribute to phenotypic instability in tumor cells

Since Illuminati events seem to be influenced by environmental stress, we sought to determine if Illuminati could contribute to malignant phenotypic instability. To test this possibility, we chose to study Illuminati in *brat* tumors. *Brat* tumors originate from type II NB-lineage intermediate progenitors that are transformed into immortal NB-like tumor stem cells (Betschinger, Mechtler, and Knoblich 2006).

To asses Illuminati frequency during *brat* tumor development we quantified GFP clones in *brat^K06028^* mutant and Ctrl flies that carried *GAL4::UAS-GFP* combined with *Tub-GAL80^Y29.7^*. We choose this *GAL80* insertion on the Y chromosome because of the ease to unambiguously test for Y chromosome loss by FISH. We could not detect significant differences in Illuminati frequency between *Tub-GAL80^Y29.7^* control *and Tub-GAL80^Y29.7^ brat* brains before larvae reached third instar stages (Figure 7A). However, the rate of Illuminati cells per lobe in *brat* brains increased dramatically afterwards and remained significantly greater than in control brains with most *Tub-GAL80^Y29.7^ brat^mut^* brains presenting more than 5 illuminati clones per brain and up 20 in extreme cases (Figure 7A). Importantly, the FISH probe against the Y chromosome confirmed that all (n=30) GFP+ clones analyzed in *Tub-GAL80^Y29.7^ brat* brains retained the Y-GAL80 chromosome (Figure 7B).

**Figure 7:**
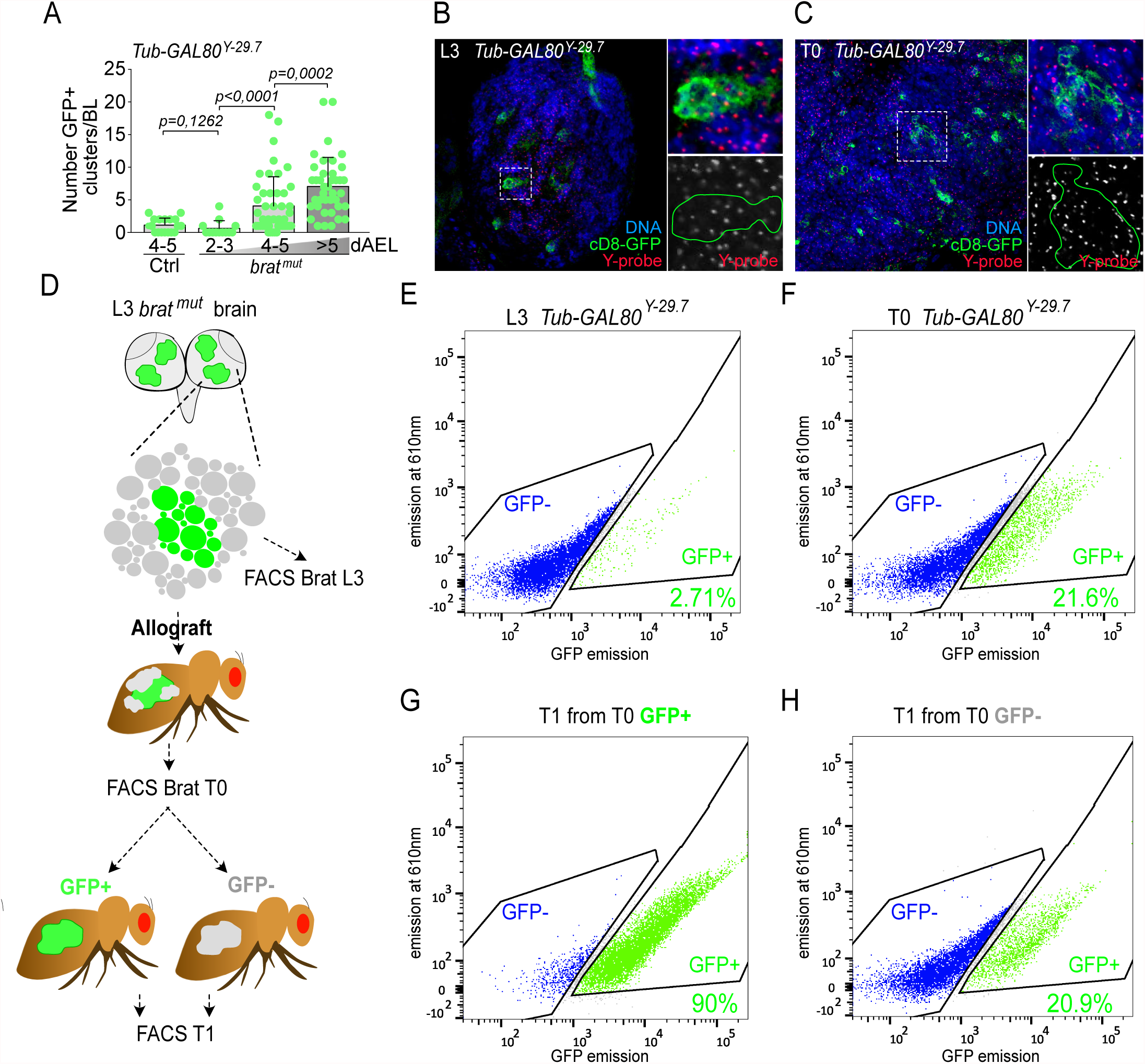
Illuminati is maintained in *brat*-induced tumors and can appear *de novo*. (A) Dot plot showing the number of illuminati clones/brain lobe in *Tub-GAL80^Y-29.7^* control and *Tub-GAL80^Y-29.7^ brat^mut^* brains in consecutive days after egg laying (dAEL). Statistical significance was determined by a Mann-Whitney test and p corresponds to the p-value. Error bars correspond to the means ± SD. (B-C) FISH with Y-probes (red and grey) of whole mount brains combined with GFP labelling (green) and DAPI for DNA (blue). Continuous dotted lines surround illuminati clones in (B) L3 brains and (C) T0 tumor after allograft. (D) Schematic representation of the protocol to analyze Illuminati behavior and stability upon tumorigenesis based on FACS analysis and successive transplantation assays. (E-H) Representative dot plot of illuminati cells quantified by FACS. (E-F) The fraction of illuminati cells massively increased from (E) before to (F) after allografting (n=2 independent experiments). (G) The rate of illuminati cells in T1 from transplantation of T0 GFP+ cells is maintained. (H) A fraction of illuminati cells emerged in T1 from transplantation of T0 GFP^_^ (n=1 experiment).

We then decided to follow Illuminati behavior during the period of massive growth that takes place upon allograft of larval brain tumors into adult hosts and to use FACS to quantify the fraction of GFP+ and GFP-cells within the tumors (Figure 7D-F). It has been estimated that from allograft to host’s death, which in most cases occurs in less than two weeks, the tumor mass expands more than a hundred-fold the original mass (Caussinus and Gonzalez 2005). Interestingly, the fraction of GFP+ cells increased massively (5-8 fold) after allografting in two independent experiments (Figure 7D-F). Once more, all the GFP+ clones analyzed by FISH in the tumor (n=70) retained the Y-GAL80 chromosome (Figure 7C). These results show that Illuminati is enhanced in *brat* tumors.

Taking advantage of the extended live of allografted *brat* tumors we decided to assess the behavior of Illuminati during this period of massive growth. To this end we allografted samples than contained only GFP+ or GFP-cells purified by FACS from a *Tub-GAL80^Y29.7^ brat* allograft at T0 (Figure 7F) and used FACS again to quantify the fraction of GFP- and GFP+ cells in the resulting T1 allograft (Figure 7D). We found that most (90.0%) of the cells in these T1 tumors that develop upon allograft of pure GFP+ T0 cells remain GFP+, thus strongly suggesting that the memory effect that stabilizes GFP expression in Illuminati NBs remains under malignant growth conditions at time points that are well beyond the constrains of normal larval development (Figure 7F-G). In addition, we found that a significant fraction (20.9%) of the cells in T1 tumors that develop upon allograft of pure GFP-T0 cells became GFP+ (Figure 7F and H), thus showing that *de novo* Illuminati events occur in T1 allografts. This time period corresponds roughly to about 3 weeks after the tumor started to develop in the larva.

Altogether, our results reveal that Illuminati is strongly enhanced during *brat* tumor growth, which in turn suggest that this new phenomenon may represent a previously unsuspected type of chromatin instability maintained in tumor cells.

## Discussion

We have generated a collection of *Drosophila* lines carrying a series of *GAL80* transgenes inserted in all major chromosomes that can be used as sensors of genome instability (GI). GI affecting Gal80 function in individuals that, in addition to *GAL80*, carry a *UAS-GFP* transgene and express *GAL4* ubiquitously generates an irreversible upregulation of GFP expression that labels the cell in which the GI event took place and its offspring. The localization and frequency of such GFP clones provide quantitated estimates of the extent of GI in different organs, tissues, and cell types, while clone size reflects the time in development when the triggering GI event occurred. Indeed, these reporters are sensitive to all GI types (e.g. CIN, CNV, point mutations, etc) and can be used to study somatic mosaicism *in vivo* with one-cell, one GI-event resolution.

All the *GAL80* lines that we generated behave as expected, efficiently repressing GFP expression in *GAL4::UAS-GFP* cells in most larval tissues. In larval brains, however, we found an unexpected number of fluorescent clones in the central brain region that in most cases included one NB and its offspring. We have named these unscheduled clones “Illuminati”. FISH and genomic DNA sequencing demonstrate that Illuminati clones retain the original wild-type copy of the *GAL80* transgene hence ruling out GI and strongly suggesting an epigenetic mechanism as the cause of Illuminati.

There are two important technical considerations to be derived from the discovery of Illuminati. The first is a note of caution regarding the use of techniques that like MARCM (T. Lee and Luo 1999) and “gypsy-trap” (W. Li et al. 2013) are based on the loss of *GAL80*. Our findings show that in larval brains, and particularly in central brain NBs, a fraction of the clones that are assumed to result from MARCM, or from *de novo* integration of gypsy elements, may actually be Illuminati clones in which recombination, or gypsy integration, have not taken place.

The second regards the potential of Illuminati as a method to generate GFP-labelled clones expressing any UAS-driven sequence of interest in central brain neuroblasts without the need to induce mitotic recombination. Importantly, taking advantage of collection of *GAL80* insertions reported here, it is possible to design experiments such that the expected number of clones can be predetermined, from many to just a few per brain. Moreover, unlike recombination-based methods that only allow for binary on/off conditions, Illuminati clones present a rather wide dynamic range of expression, which can be read out as fluorescence intensity, that may reveal dosage-dependent effects that would pass unnoticed in conventional clones.

There are interesting similitudes and differences between Illuminati and the other two types of variegated expression known in *Drosophila*: PEV and Gal4::UAS variegation. In all three cases variegation appears as clones founded by an individual cell that underwent an epigenetic shift in gene expression that was fixed and maintained through the successive cell cycles that originated the clone. Also like Gal4::UAS variegation, but unlike PEV, Illuminati does not result from proximity to heterochromatin. Moreover, certain of the PEV and Gal4::UAS modifiers that we have tested do not affect Illuminati at all, while others affect Illuminati in ways that are consistent with their effect on PEV and Gal4::UAS variegation. However, they do so to a rather low extent. Also relevant is the fact that unlike Gal4::UAS variegation that acts mostly in cis (Skora and Spradling 2010), Illuminati acts in trans, although such a difference may at least partially be due to the fact that Illuminati results from changes in Gal80 that in turn affect Gal4::UAS, rather than from changes in Gal4 itself, as it is the case in Gal4::UAS variegation.

The discovery of Illuminati identifies a previously unappreciated form of gene expression plasticity. Illuminati occurs during normal development in *Drosophila* neural stem cells and is strongly enhanced in at least one type of neural stem cell-derived malignant neoplastic tumor. During normal stem cell development, Illuminati may reflect the lack of full epigenetic competence that characterizes stem and early progenitor cells. This may contribute to make Illuminati NBs less able, than more differentiated cells, to accurately transmit non-genetic information to their progeny. Importantly, this is the case during follicle cell development and it has been proposed as a possible mechanism generating NB competence to produce different types of neurons in the *Drosophila* brain (Pearson and Doe 2003; Cleary and Doe 2006). During tumor growth, Illuminati may be a major contributor to non-genetic variability that may have a significant role in malignant transformation.

Like Gal4::UAS variegation in the ovary follicle, we expect Illuminati to provide an unprecedented means towards understanding how alterations in the chromatin-based machinery of epigenetic inheritance contribute to neural stem cell development under normal and disease conditions. It will be interesting to determine if Illuminati also occurs in the brain of other animals.

## Acknowledgments

We thank F., S. and L. Scaerou for numerous discussions concerning the name of the phenomenon described here culminating with the final suggestion of Illuminati. We acknowledge the PiCT-IBiSA platform and Nikon Imaging Center at Institut Curie microscopy. We thank V. Marthiens, S. Gemble, M. Budzyk, F. Leulier, J. Merlet, P. Tran, G. Almouzni, the Bardin team and members of the Gonzalez laboratory for helpful discussion and/or comments on the manuscript. This work was supported by (ERC) AdG 2011 294603, ERC CoG (ChromoNumber 725907), Spanish MINECO PGC2018-097372-B-100 funded by European Regional Development Fund (ERDF)/Ministry of Science, Innovation and Universities-Spanish State Research Agency, Institut Curie and the CNRS. A.G was funded by FRM (ECO20170637529) and LLCC (IP/SC-16533) fellowships. The Basto lab is a member of the Labex Cell(n)Scale.

## Materials and Methods

### Fly husbandry and fly stocks

For most experiments, flies were raised in plastic vials containing homemade standard *Drosophila* rich culture medium (0.75% agar, 3.5% organic wheat flour, 5% yeast, 5.5% sugar, 2.5% nipagin, 1% penicillin-streptomycin (Gibco #15140), and 0.4% propanic acid). Fly stocks were maintained at 22°C and experimental crosses at 22°C or 25°C. For food restriction experiment, flies were raised on homemade protein-poor medium (0.75% agar, 7% cornmeal, 1.4% yeast, 5.2% sugar, 1.4% nipagin) at 22°C and compared to flies raised on homemade standard rich medium at 22°C. For temperature variation experiments, flies were laying eggs for 24hours and tubes containing progeny were maintained at 18°C, 22°C, 25°C or 29°C for 7, 5, 5 or 4 days, prior dissection, respectively. For MARCM experiments to estimate proliferation in wing discs and brain lobes, females of the genotype *hsFLP Tub-GAL80^X-1C2^ neoFRT19A* were crossed to *neoFRT19A* males; crosses were kept at 25°C, heat-shocked for 1.5h at 37°C, 48-72h AEL, and brains and wing discs of female larvae were dissected at 96-120h AEL. For sas4 MARCM experiments, fly crosses were kept at 22°C. L2 progenies were heat-shocked 1hour at 37°C in a water bath and maintained at 22°C for 48±12 hours before dissection.

### Tub-GAL80 Drosophila lines establishment

ΦC31-recombination attB P[acman] system for *Tub-GAL80* insertions on chromosome X, II and III: the plasmid containing codon-optimized *GAL80* sequence driven by a tubulin promoter is a gift from Allison Bardin and corresponds to the combination of *pattB-tubP-SV40* - generated by Lee and Luo (T. Lee and Luo 1999)- with the codon optimized *GAL80* sequence from *pBPGAL80Uw-6* - a gift from Gerald Rubin (Addgene plasmid #26236, http://n2t.net/addgene:26236; RRID:Addgene_26236) (Pfeiffer et al. 2010). Then, the plasmid was inserted in a P[acman] vector and send to Bestgene® company to integrate it into the *Drosophila* genome at specific insertion sites using PhiC31 integrase-mediated transgenesis system.

To obtain *Drosophila* recombinants carrying two copies of *Tub-GAL80* on the same chromosome, we used female meiotic recombination and selected recombination events based on the fly eyes colour, a method widely used to generate *Drosophila* recombinants. Indeed, all *Tub-GAL80* are associated with the white^+^ transgene expressing marker, which is used as a marker for efficient transgene insertion as it confers yellow to red eye colour. Simply, in the presence of two copies, as for efficient recombination, fly eyes display a strong red colour.

The *P{w^+^, tubP-GAL80}LL1* transgene terminally located on the X chromosome (T. Lee and Luo 1999), was recombined onto a *y w f* marked *C(1;Y)* chromosome. The resulting *C(1;Y) P{w+, tubP-GAL80}LL1 w f* chromosome was mutagenized with 4000-R of X-rays to induce deletions that remove most of the X chromosome.

### Deep sequencing

Illumina reads were aligned by BWA software (version 0.7.10; (H. Li and Durbin 2009)) using default options and data analysed using SAM tools (version 0.1.19; (H. Li et al. 2009)). Whole-genome alignment were performed using the drosophila melanogaster release 6.02 as reference. (http://flybase.org/static_pages/docs/release_notes.html).

### Immunofluorescence of *Drosophila* larval whole mount tissues

Wandering third-instar larval (L3) brains and imaginal discs were dissected in fresh Phosphate-Buffered Saline 10X (PBS, VWR #L182-10) and fixed for 30 minutes (min) at room temperature (RT) in 4% paraformaldehyde (EMS # 15710) diluted in PBS. Fixed tissues were washed and permeabilized three times 15 min in PBST3 or PBST1 (PBS, 0,3% or 0,1% Triton X-100, Euromedex #2000-C). For antibody staining, larval tissues were incubated in primary antibodies diluted in PBST3 or PBST1 overnight at 4°C in a humid chamber. After 3×15 min washes in PBST3 or PBST1, tissues were incubated in secondary antibodies diluted in PBST3 or PBST1, O/N at 4°C and protected from light in a humid chamber. Tissues were then washed 3×15 min in PBST3 or PBST1, rinsed in PBS and mounted between slides (Thermo Fisher Scientific #AA00008232E00MNT10) and 12-mm circular cover glasses (Marienfield Superior #0111520) with 5µl of homemade mounting medium (1,25% n-propyl gallate, 75% glycerol, 25% H_2_O).

For GFP labeling of larval brains and imaginal discs, two different protocols were used: 1) one step of O/N incubation at 4°C with GFP booster (1:250, Alexa Fluor ® 488 Chromotek #gb2AF488), Phalloidin-647 (1:250, Thermo Fisher Scientific #A-22287) and DAPI (1:1000, Thermo Fisher Scientific #62248) were performed followed by 3×15 min washes in PBST3, a rinse in PBS and mounting. 2) Brains and imaginal discs of wandering L3 larvae were dissected in fresh PBS and fixed for 20 min at RT in 4% paraformaldehyde diluted in PBS. Fixed tissues were washed three times for 10 min in PBS, permeabilized 1h in PBST3, washed for 10 min in PBS and stained with DAPI (1:1000) for 20 min, washed again 10 min in PBS and mounted in Vectashield.

Primary antibodies used in this study are: chicken (chk) anti-GFP (1:1000, Abcam #ab13970), guinea pig (GP) anti-Deadpan (Dpn) (1:1000, J. Skeath), mouse anti-Prospero (1:500, MR1A, DSHB), rat anti-Elav (1:100, 7EA10, DSHB), mouse anti-Repo (1:500, 8D15, DSHB), rabbit (Rb) anti-Sas4 (1:500, Basto et al 2006), GP anti-Centrosomin (Cnn) (1:1000, E. Lucas and J.W.R.).

Secondary antibodies (1:250) used in this study are: chk-488 (Thermo Fisher Scientific #A-11039), Rat-546 (Thermo Fisher Scientific #A-11081), Rb-568 (Thermo Fisher Scientific #A-10042), mouse-546 (Thermo Fisher Scientific #A-11030) and GP-647 (Thermo Fisher Scientific #A-21450).

Images were acquired with 40x (NA 1.25), 63x (NA 1.32) or 100x (NA 1.4) oil objectives on a wide-field Inverted Spinning Disk Confocal Gattaca/Nikon (a Yokagawa CSU-W1 spinning head mounted on a Nikon Ti-E inverted microscope equipped with a camera complementary metal-oxide semiconductor 1.200 x 1.200 Prime95B; Photometrics). Intervals for z-stack acquisitions were set to 0.5 to 1.5µm using Metamorph software.

### Live imaging of *Drosophila* larval brains

Mid second-instar larval (L2) brains were dissected in Schneider’s *Drosophila* medium (Gibco #21720024) supplemented with 10% heat-inactivated fetal bovine serum (Gibco #10500), penicillin (100 U/ml) and streptomycin (100 µg/ml). Several brains were placed in 10µl of medium on a glass-bottom dish (Dutcher #627870), covered with a permeable membrane (Standard YSI), and sealed around the membrane borders with oil 10 S Voltalef (VWR Chemicals). Images were acquired with 60x oil objective (NA 1.4) on two microscopes: an Inverted Spinning Disk Confocal Roper/Nikon (a Yokagawa CSU-X1 spinning head mounted on a Nikon Ti-E inverted microscope equipped with a camera EMCCD 512 x 512 Evolve; Photometrics) and the wide-field Inverted Spinning Disk Confocal Gattaca/Nikon (a Yokagawa CSU-W1 spinning head mounted on a Nikon Ti-E inverted microscope equipped with a camera complementary metal-oxide semiconductor 1.200 x 1.200 Prime95B; Photometrics), controlled by Metamorph software. For both microscopes, images were acquired at time intervals spanning 30 min and 50 z-stacks of 1.5 µm.

### DNA Fluorescent *in situ* hybridization

After fixation, permeabilization and O/N incubation with GFP booster (description below), brains were washed 3×15min in PBT3 and fixed a second time 30 min in 4% PFA. Then, brains were rinse 3x in PBS, washed 1×5 min in 2xSSCT (2X Saline Sodium Citrate (Euromedex #EU0300-A) 0,1% Tween-20 (Sigma Aldrich #P1379) diluted in water), 1×5 min in 2xSSCT/50% formamide (Sigma Aldrich #47671), transferred in pre-warmed 2xSSCT/50% formamide and pre-hybridized 3 min at 92°C. In the meantime, DNA probes diluted in the Hybridization Buffer (20%dextransulfate (Sigma Aldrich #D8906), 2xSSCT, 50%formamide, 0,5mg/ml salmon DNA sperm (Sigma Aldrich #D1626)) were denature at 92°C. After removal of the supernatant, brains were incubated in the probes solutions and hybridize 5 min at 92°C and O/N at 37°C. Brains were then rinse at RT, washed 1×10 min at 60°C and 1×5 min at RT in 2xSSCT. Finally, after a rinse in PBS brains were mounted as described below. DNA probes used in this study were against chromosomes X (80ng/µl), II (40ng/µl) and III (80ng/µl). FISH for the Y chromosome was performed with a probe to detect the AATAC repeat, one of the Y-specific satellites of the *Drosophila* genome (Bonaccorsi and Lohe 1991). Chromosome spreads of mitotic chromosomes (karyotypes) were prepared and stained with DAPI following standard procedure (Fanti and Pimpinelli 2004). Hybridisation conditions were as described in (Dernburg 2011) (http://cshprotocols.cshlp.org/content/2011/12/pdb.prot066902.abstract).

### RNA Fluorescent *in situ* hybridization

L3 brains were dissected in fresh PBS, fixed 30 min in 4% formaldehyde (EMS # 15686) and washed and permeabilized in PBSTw (PBS 0,3% Tween-20). Brains were incubated with GFP booster and Phalloidin diluted in PBSTw, O/N at 4°C in a humid chamber. After 3×15 min washes PBSTw, RNA hybridization was performed as described by Yang and colleagues (Yang et al. 2017). RNA probes against *GAL80* were designed by the biosearchtech® technical support team (https://www.biosearchtech.com/) and labeled with quasar 570.

### Allografts

Allografts were performed as described in (Rossi and Gonzalez 2015).

### Generation of Brat tumors in illuminati clones

Males of the genotype *y^1^ sc* v^1^ sev^1^; UAS-brat-RNAi* (P{TRiP.HMS01121}attP2) (Ni et al. 2011) were crossed to females *Tub-GAL80^X-1C2^; Tub-GAL4 UAS-cD8::GFP/CyO, S, Tb* and incubated at 29°C. Five day old female larvae of the genotype *Tub-GAL80^X-1C2^/ +; Tub-GAL4 UAS-cD8::GFP/+; UAS-brat-RNAi/ +* were dissected and brain lobes were allografted into RFP-α-Tub female hosts. Tumors were cultured 12-13 days at 29°C, flies dissected and GFP+ tumor cells were prepared for DNA extraction.

### DNA extraction, PCR and sequencing

Genomic DNA was extracted from tumors using standard Phenol/Chlorophorm protocol and precipitated with Isopropanol in the presence of Pellet Paint (Merck). The amplification was done using 2 ng of DNA and KOD hot star (MercK). PCR fragments were sequenced by EurofinsGenomics. Primers are listed below.

### PCR primer

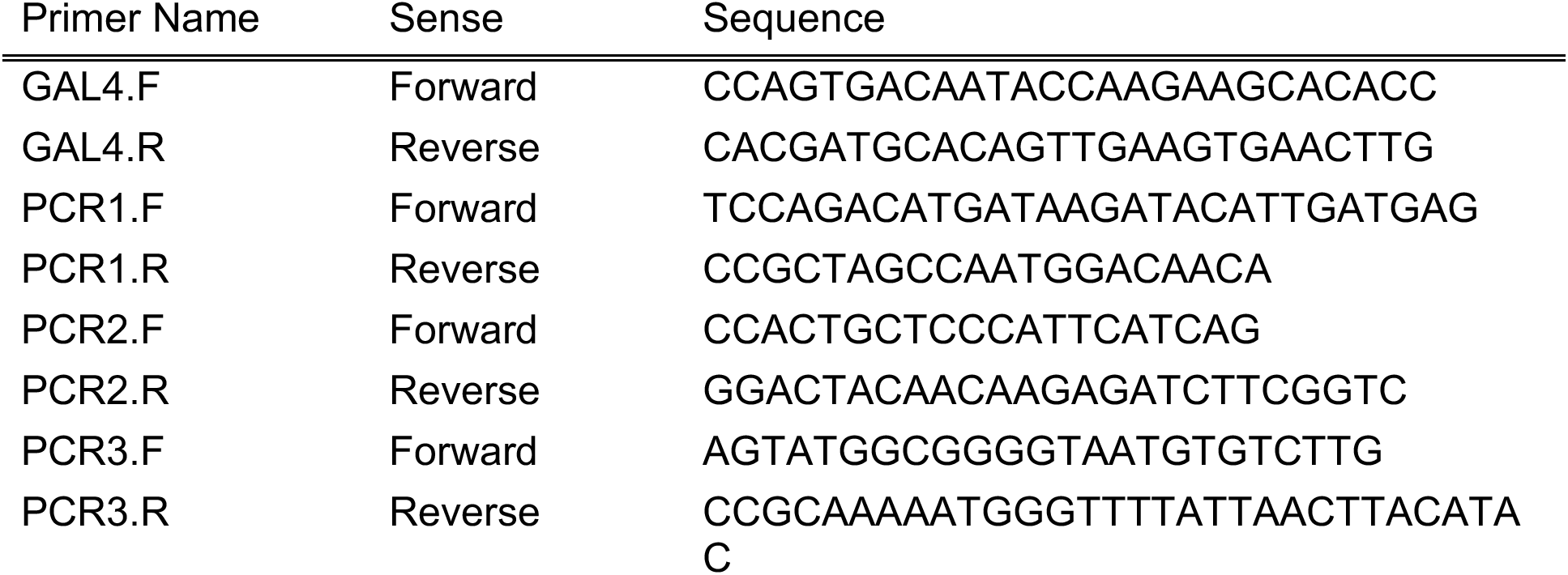

### Sequencing primer

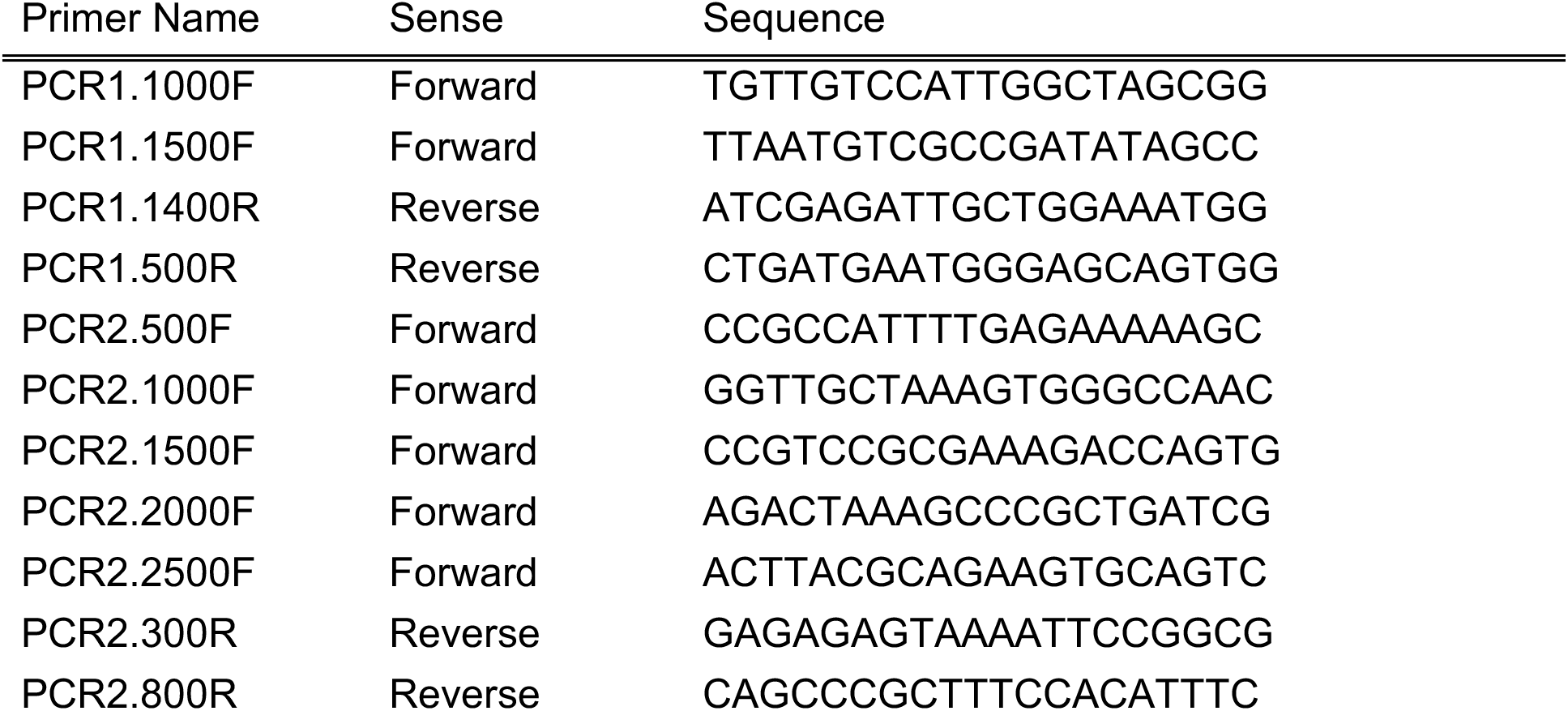
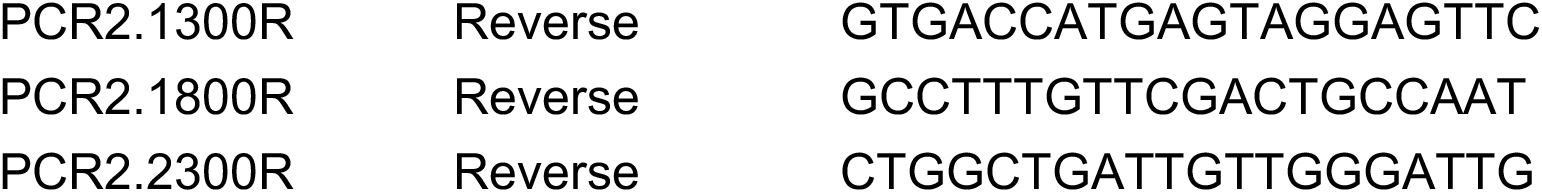

### FACS

FACS was performed as in (Castellanos, Dominguez, and Gonzalez 2008).

---

**Supplementary Figure 1:**
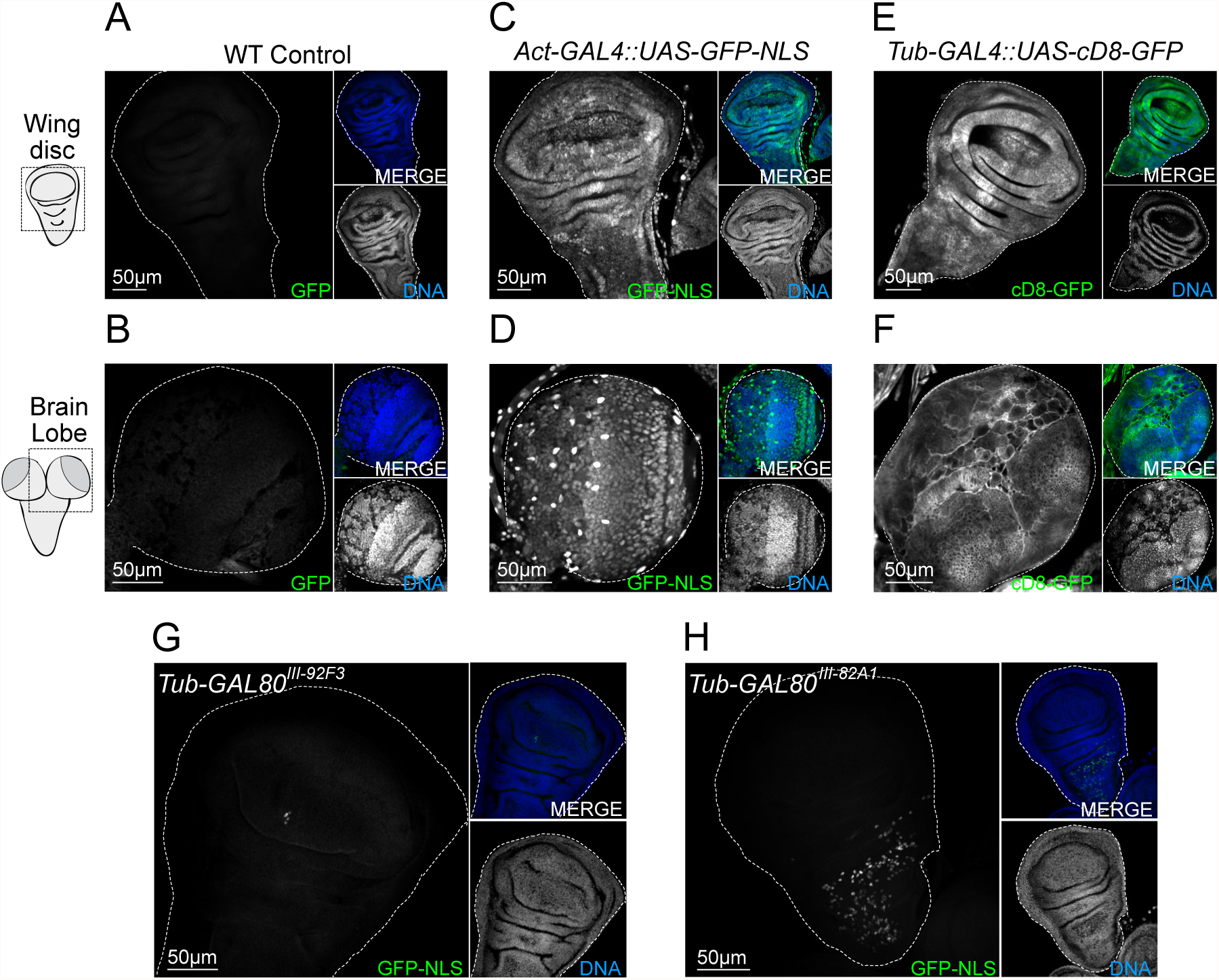
Analysis of Control and *Tub-GAL80* wing discs and brains. (A-H) Images of whole mount tissues labelled for GFP (grey and green in large and small insets respectively) and DNA (blue and grey in small insets) in WT (A) wing disc and (B) brain lobe, Act-GAL4::UAS-GFP-NLS (C) wing disc and (D) brain lobe, *Tub-GAL4::-UAS-cD8-GFP* (E) wing disc and (F) brain lobe and wing discs from (G) Tub-GAL80^III92F3^ and (H) Tub-GAL80^III82A1^. White dotted lines surround tissues.

**Supplementary Figure 2:**
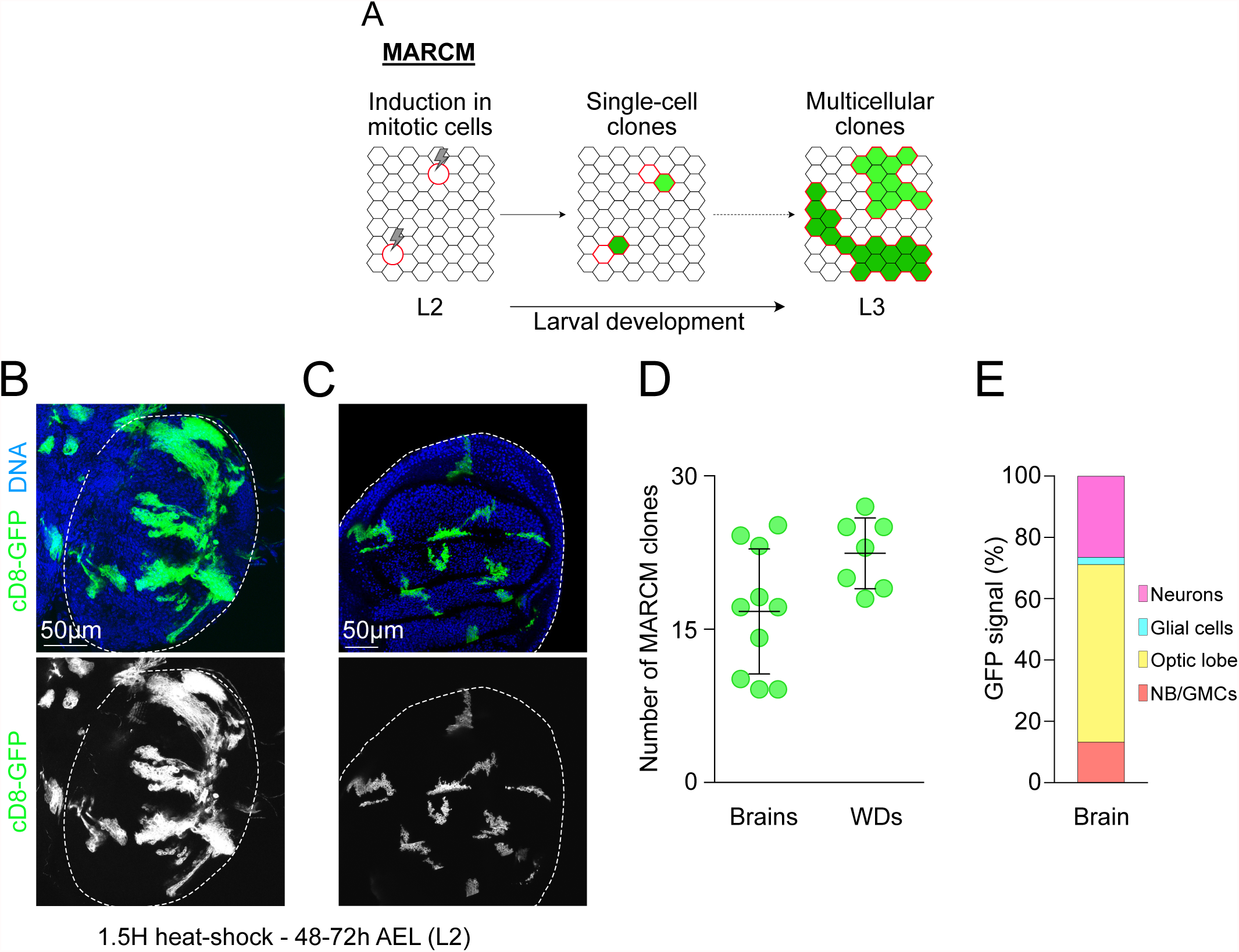
The presence of high level of GFP+ cells in the brain is not explain by a higher proliferation rate. (A) Schematic representation of the clone induction with MARCM system during development as a proxy to determine the frequency of mitosis and proliferation in tissues. (B-C) Images of MARCM clones in whole mount brain lobe (B) and wing disc labeled for GFP (grey and green) and with DAPI for DNA (blue). White dotted lines surround tissues. (D) Dot plot showing the number of MARCM clones in brain lobes (n=10 BLs) and wing discs (n=7 Wing discs). Error bars correspond to the means ± SD. (E) Graph bar showing the percentage of cells showing GFP+ signals for each cell type of the brain lobe: NB/GMCs clusters (red), cells from the optic lobe (yellow), individual glial cells (cyan blue) and individual neurons (pink).

**Supplementary Figure 3:**
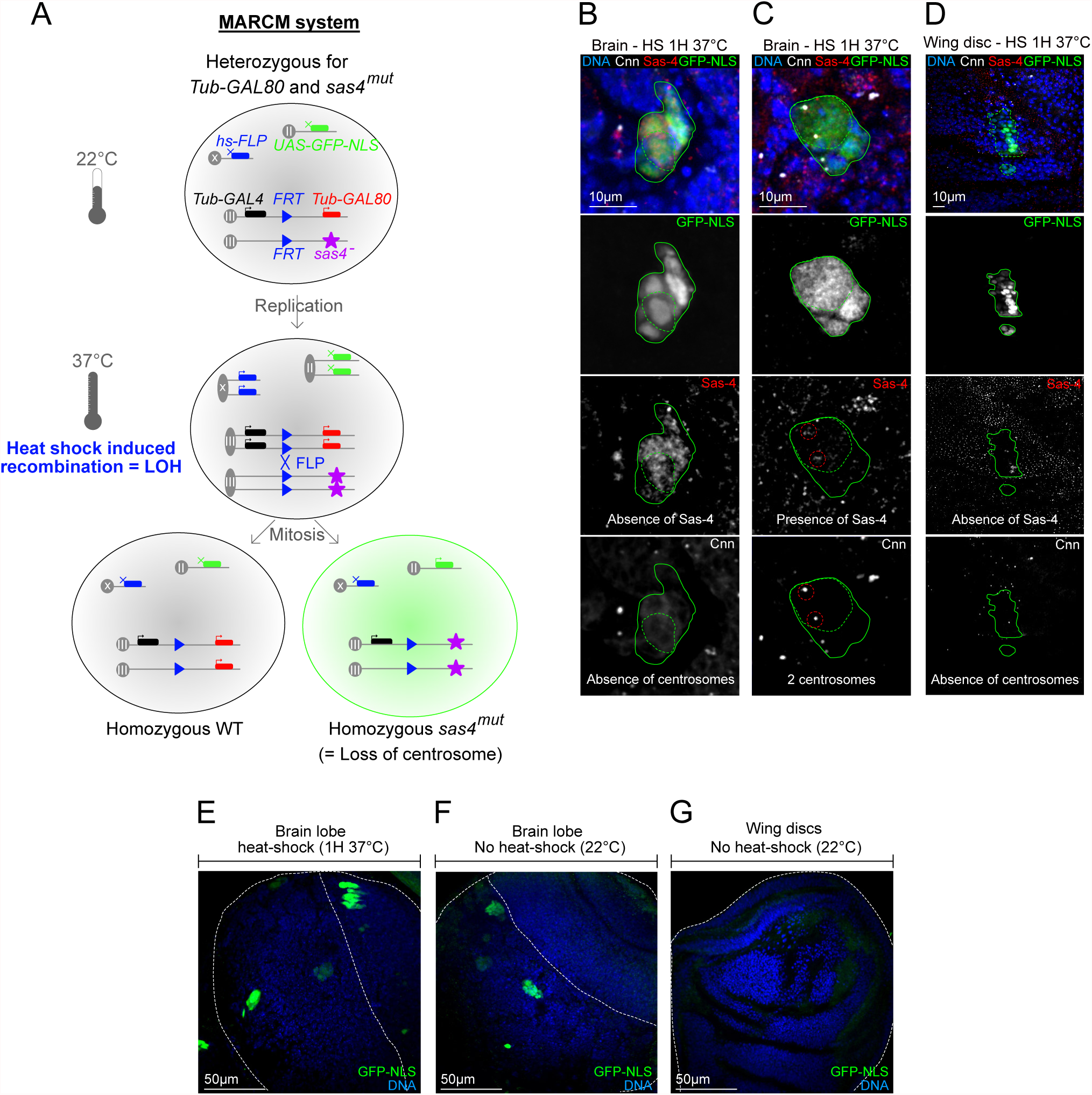
Analysis of Tub-GAL80 lines alerts its use for MARCM analysis. (A) Schematic representation of the MARCM system to induce mutant clones labeled. After recombination of FRT sites by the heat-induced FLP recombinase, the daughter cells lose heterozygosity. One cell becomes homozygous *sas4^mut^* and labeled with GFP due to the concomitant loss of the *Tub-GAL80* sequence. The other cell becomes homozygous WT and it is unlabeled. (B-D) Images of GFP+ clones in (B-C) brain lobes and (D) wing disc of *hs-FLP/+, UAS-GFP-NLS/+; Tub-GAL4,FRT82B,Tub-GAL80/FRT82B,sas4^mut^* flies heat-shocked at 37°C for 1 hour and labeled with antibodies against GFP (green and grey), Sas-4 (red and grey), Cnn (grey) and with DAPI for DNA (blue). Green continuous and dotted lines surround GFP+ clones and NBs, respectively. (B) *Sas4* mutant GFP+ NB without centrosomes. (C) Wild type GFP+ NB with two centrosomes. (D) *Sas4* mutant GFP+ clones in the WD. (E-G) Images of whole mount (E-F) brain lobes and (G) wing discs labeled for GFP (green) and with DAPI for DNA (blue). (E) Presence of GFP+ clones in brain lobes after 1 hour of heat-shocked at 37°C. (F-G) In the absence of heat-shock, (F) brain lobes present GFP+ clones, in contrast to (G) wing discs that are GFP^_^.

**Supplementary Figure 4:**
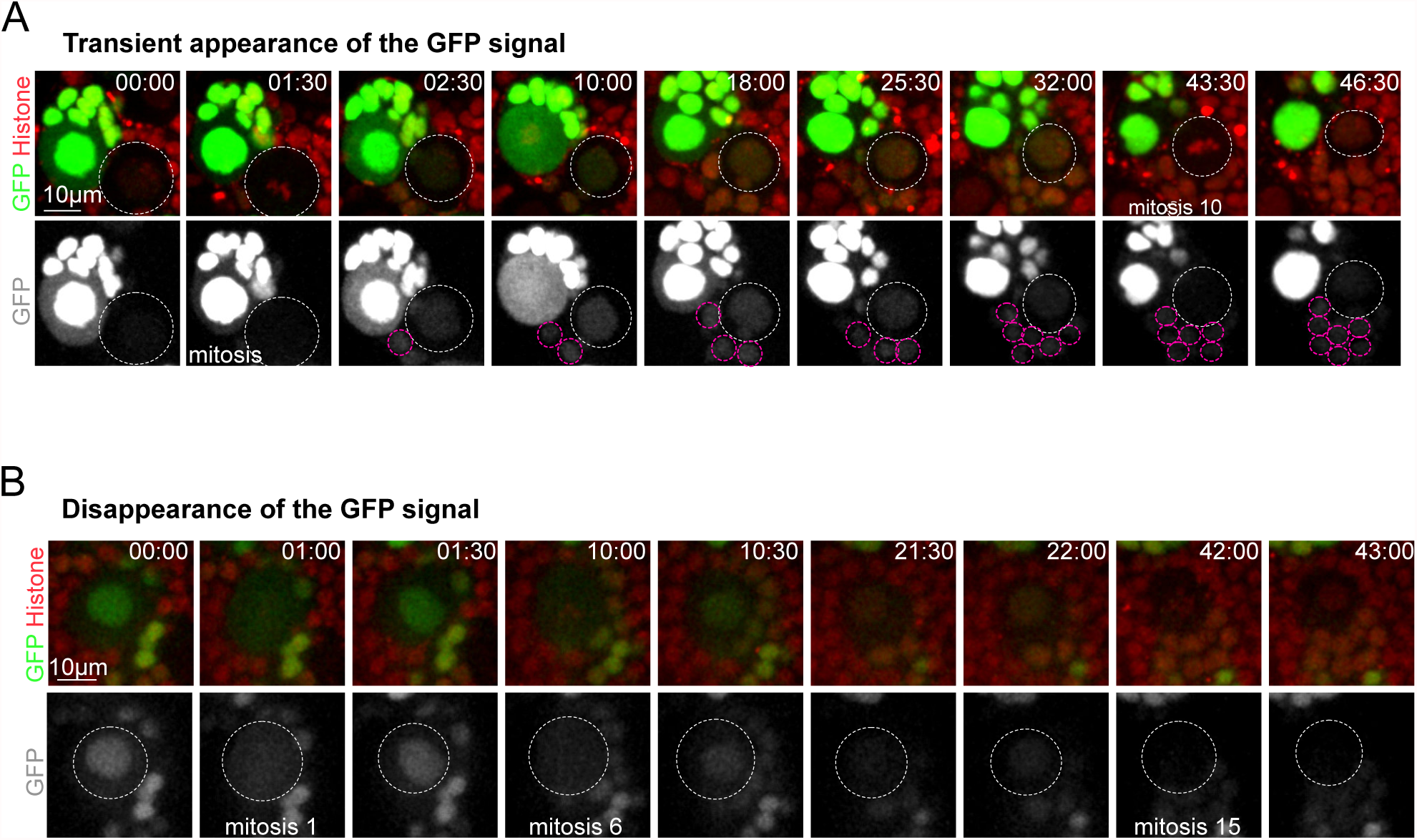
Illuminati expression is dynamic and reversible in extremely rare cases. (A-B) Stills of time-lapse movies of mitotic NBs expressing *2xTub-GAL80^X-5B8,19E7^, GAL4::GFP-NLS* (green) and *histone-RFP* (red) to monitor GFP and chromosome dynamics. White and pink dotted circles surround NBs and daughter GMCs, respectively. GFP signal is dynamic in rare cases: (A) the transient appearance or (A) the disappearance of the GFP signal.

**Supplementary Figure 5:**
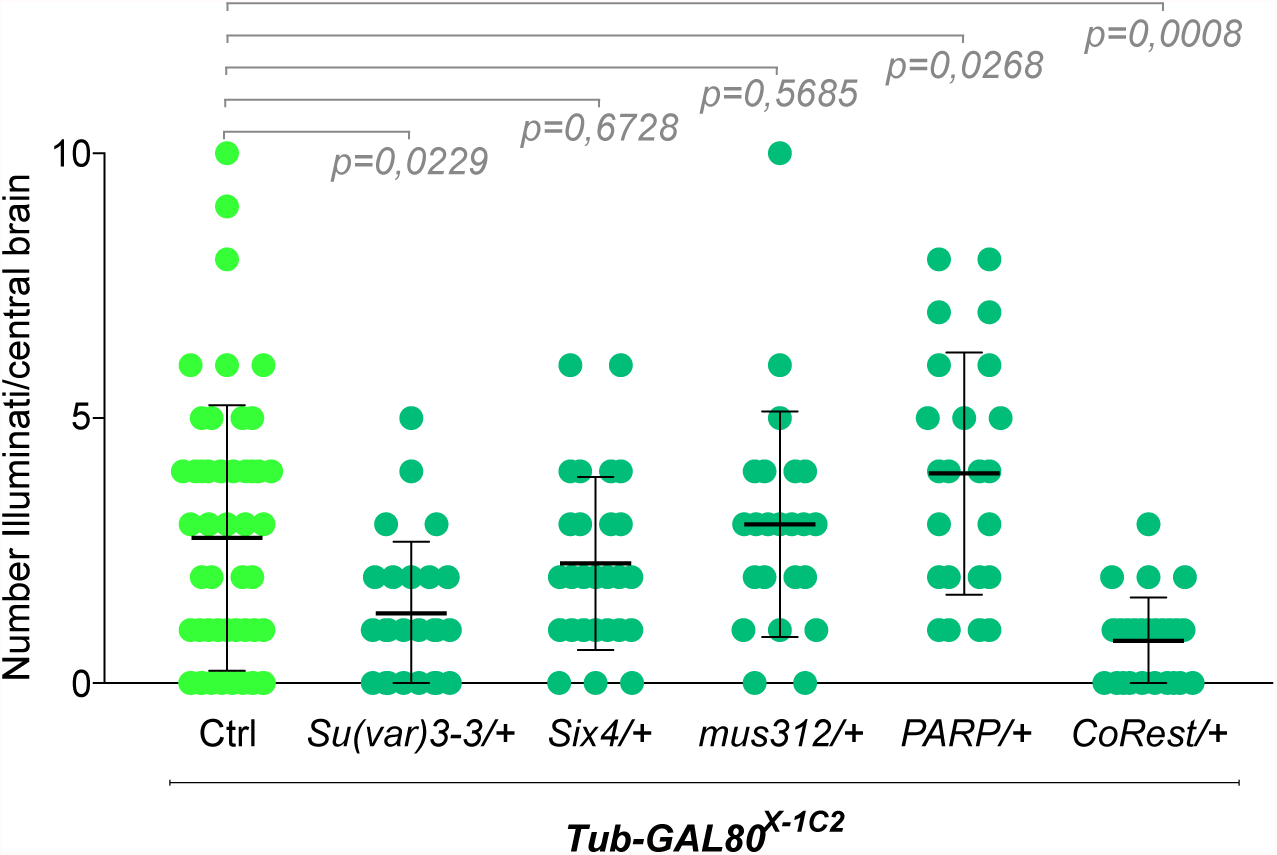
PEV and Gal4::UAS variegation do not explain the lack in *GAL80* expression. Dot plot showing the number of Illuminati/central brain in *Tub-GAL80^X-1C2^* Ctrl (n=52 BLs) and in heterozygous mutant for *Su(var)3-3* (n=26 BLs), *six4* (n=28 BLs), *mus312* (n=24 BLs), *PARP* (n=24 BLs) and *CoRest* (n=26 BLs). Statistical significance was determined by a Mann-Whitney test and p corresponds to the p-value. Error bars correspond to the means ± SD.

## Notes

### Competing Interest Statement

The authors have declared no competing interest.

